# Intrinsic microtubule destabilization of multiciliated choroid plexus epithelial cells during postnatal lifetime

**DOI:** 10.1101/2023.01.10.523428

**Authors:** Kim Hoa Ho, Valentina Scarpetta, Chiara Salio, Elisa D’Este, Martin Meschkat, Christian A. Wurm, Matthias Kneussel, Carsten Janke, Maria M. Magiera, Marco Sassoè-Pognetto, Monika S. Brill, Annarita Patrizi

**Affiliations:** Schaller Research Group, German Cancer Research Center (DKFZ), Heidelberg, Germany; Faculty of Biosciences, Heidelberg University, Heidelberg, Germany; Department of Neurosciences “Rita Levi Montalcini”, University of Turin, Turin, Italy; Department of Veterinary Sciences, University of Turin, Grugliasco, Italy; Optical Microscopy Facility, Max-Planck-Institute for Medical Research, Heidelberg, Germany; Abberior Instruments GmbH, Göttingen, Germany; Department of Molecular Neurogenetics, Center for Molecular Neurobiology, ZMNH, University Medical Center Hamburg-Eppendorf, Hamburg, Germany; Institut Curie, PSL Research University, CNRS UMR3348, Orsay, France; Université Paris-Saclay, CNRS UMR3348, Orsay, France; Institute of Neuronal Cell Biology, Technical University of Munich, Munich, Germany; Munich Cluster of Systems Neurology (SyNergy), Munich, Germany; Interdisciplinary Center for Neuroscience, Heidelberg University, Heidelberg, Germany; Zentrum für Molekulare Biologie der Universität Heidelberg, DKFZ-ZMBH Alliance, Heidelberg, Germany

## Abstract

Choroid plexus (ChP) epithelium is composed of specialized multiciliated cells. By using multiple microscopic techniques, biochemical approaches in various mutant mice and longitudinal analysis from mouse embryogenesis to aging, we show that ChP cilia are built on a gradient of events which are spatio-temporally regulated. We uncover that ChP cilia develop prenatally since early tissue morphogenesis, and proceeds as a multi-step process characterized by basal body multiplication and axoneme formation directly at the apical cellular compartment. Our data also show that choroid plexus cilia contain both primary and motile features. Remarkably, we demonstrate that ChP cilia undergo axoneme resorption, starting from early youth, through a tubulin destabilization process, which is primarily controlled by polyglutamylation levels and could be mitigated by the removal of the microtubule-severing enzyme spastin. Notably, we demonstrate that this phenotype is preserved in human samples.

## Main

Choroid plexuses (ChPs) are specialized endothelial-epithelial barriers located throughout the ventricular system of the brain. The ChP is responsible for the synthesis and secretion of cerebrospinal fluid (CSF), controlling the homeostasis of the central nervous system (CNS)^1^. Recent studies have shown that ChP can sense peripheral stimuli and adapt its cellular responses accordingly^2,3^, suggesting that a major role of this organ is to provide an interface of continuous exchange between circulating blood and CSF.

The ChP arises prenatally, right after the closure of the neural tube, and individual ChPs, telencephalic (tChP) and hindbrain (hChP), emerge independently from the roof plate^4^. Most of the available papers on the development of the ChP are detailed morphometric analyses describing the progression of the ventricular system. It was shown in mammals that the majority of the dividing cells are mainly concentrated at the root of the plexus and that the tChP development is characterized by multiple histological stages^5^. It is, therefore, critical to expand our knowledge and understanding of these initial stages and to identify quantifiable key features related to the development and the maintenance of the ChP.

Choroid plexus epithelial cells are derivatives of neuroepithelial progenitors and, like the neighboring ependymal cells, are multiciliated^6^. Early electron microscopic analyses described the presence of 9+2 axonemes in cilia of several species, including humans^7^ and pigs^8^. However, most of these studies classified the ChP as part of the ependymal layer and therefore did not appreciate the potential differences between these two cell types. More recent studies confirmed that ChP cilia have a 9+2 conformation in zebrafish^9^, but showed multiple configurations, ranging from 9+0 to 9+1 and 9+2 in murine ChP^10^, implying that ChP cilia are not the same as those present in ependymal cells, and suggesting that ciliogenesis might be differently regulated in the two cell types.

Cilia are mainly composed of organized microtubules, assembled from dimers of α- and β-tubulin components^11^. These organelles are essential for both embryonic and postnatal organ development and function. Mutations in over 200 cilia genes, including tubulin genes, like *TUBB4B*^*12,13*^ and *TUBB8*^*14*^, result in a group of disorders called ciliopathies^15^. Tubulin assembly and disassembly depend on tubulin posttranslational modification (PTM) events^11^, such as acetylation, glutamylation and glycylation. Glutamylation, the reversible addition of a single or multiple glutamates to the tubulin C-terminal tails, is the most abundant tubulin modification in the mammalian brain, occurring mainly during postnatal brain maturation^16,17,18,19^. The addition and removal of glutamate residues are regulated processes which depend on selective enzymes, such as tubulin-tyrosine ligase-like (TTLL) family and deglutamylase cytosolic carboxypeptidase-like proteins (CCP), respectively^20^. The number of glutamates on tubulin tails can also promote the enzymatic severing of microtubules by spastin^21, 22^ and katanin^23^.

To understand cilia development in ChP epithelial cells, we utilized multiple microscopic techniques ranging from confocal and super-resolution (STED) optical microscopy to serial transmission electron microscopy (TEM). We found that embryonic ChP multiciliogenesis is a multi-step event occurring along a well-defined spatio-temporal trajectory, in an unconventional, not previously described process. Unexpectedly, following the maturational trajectory of the cilia apparatus until senescence, we found that axonemes gradually disappear after weaning, leaving the majority of the ChP epithelial cells with basal bodies retreating from the cell membrane to a deeper cytoplasmic compartment. Analysis of postmortem human ChP showed a similar phenomenon, suggesting that axoneme resorption is not a species-specific occurrence. We then elucidated the mechanisms underlying axoneme resorption and revealed that the reduction of microtubule polyglutamylation is a driving event that is partially mediated by the selective microtubule-severing enzyme spastin.

## Results

### Ciliogenesis as a spatial and temporal footprint of choroid plexus epithelial cell development and maturation

Early in embryonic development, ChP epithelial cells undergo a spatially and temporally regulated program of differentiation^24,25,26^. To investigate distinct stages of epithelial cell development, we correlated cell differentiation with ciliogenesis by mapping the spatial organization with multiple markers: Ki-67 for cell proliferation, E-Cadherin for epithelial cell differentiation, and γ-tubulin (γ-Tub) and acetylated α-tubulin (AcTub), for labeling respectively the basal bodies and the cilium axonemes (Fig. 1). During early ChP morphogenesis - embryonic day (E)11.5 for hChP and E12.5 for tChP^27^ - nearly all cells were proliferating, as shown by labeling for Ki-67. In contrast, E-Cadherin signal was barely detectable (Extended data Fig. 1a). At this stage, immunostaining for cilia showed numerous basal bodies but no axoneme along the whole ChP length (Extended data Fig. 1b). Taken together, these results suggest that at this early stage the ChP consists of proliferative progenitors that have not yet acquired the markers of ciliation, indicating that ChP epithelial cells may still have progenitor capacity.

**Figure 1.**
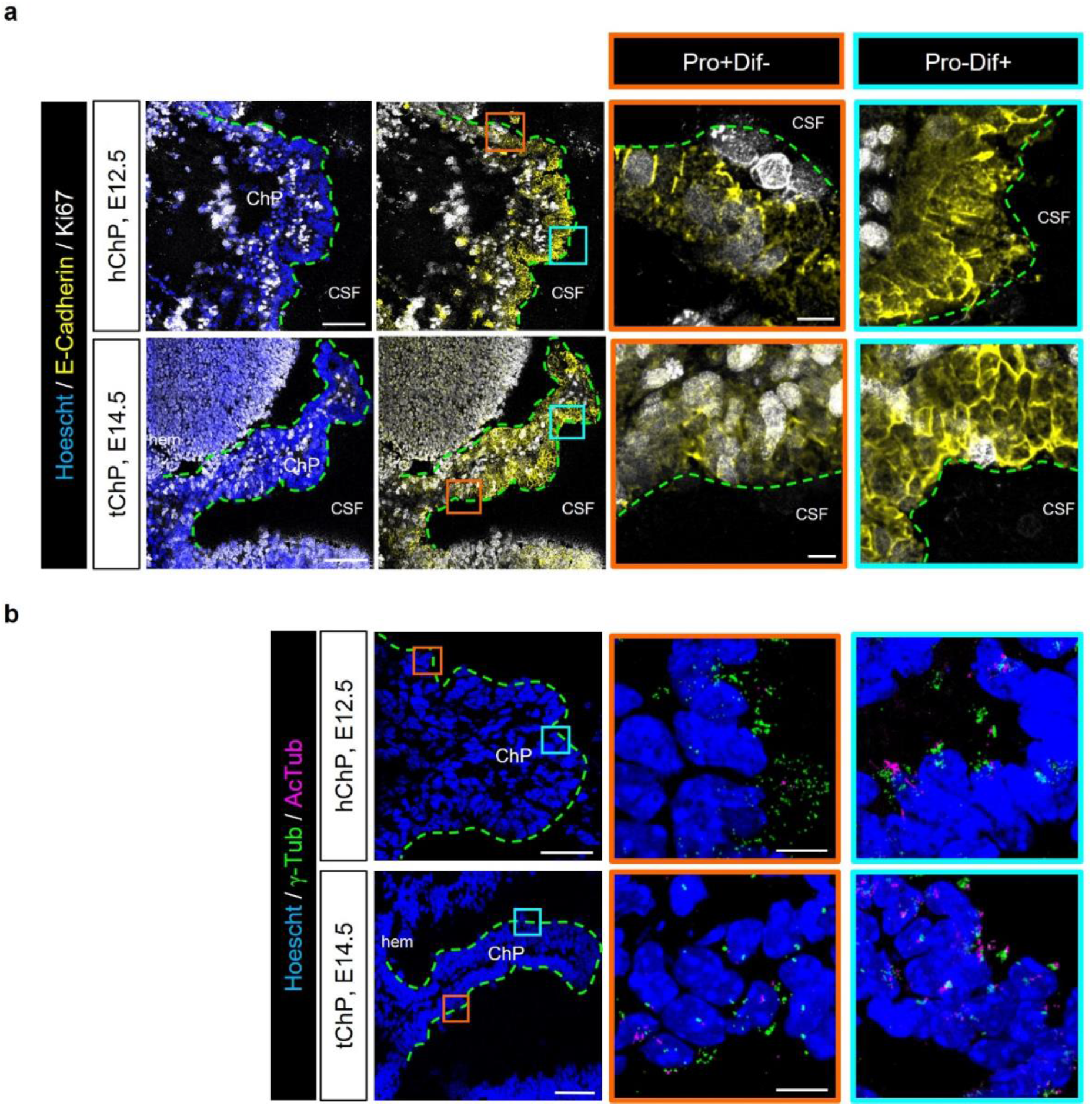
Ciliogenesis in choroid plexus epithelial cells correlates with cellular differentiation. **a, b** Immunofluorescence images of hindbrain (h) and telencephalic (t) choroid plexus (ChP) at E12.5 and E14.5, respectively. **a**, Sagittal brain sections were stained with antibodies against E-cadherin (yellow) and Ki67 (white). Orange boxes feature ChP proximate to the ventricular wall (hem) where epithelial choroid plexus cells are Ki67-positive and E-cadherin-negative (Prof+Dif-). Cyan boxes feature ChP distant from the ventricular wall where choroid plexus epithelial cells are E-cadherin-positive and Ki67-negative (Prof-Dif+). CSF: cerebrospinal fluid. **b**, Sagittal brain sections were stained with antibodies against γ-Tubulin (γ-Tub, green) and acetylated tubulin (AcTub, magenta). Orange and cyan boxes feature ChP proximate to or distant from the ventricular wall (hem), respectively. Scale bars: overview: 100 μm, inset: 10 μm.

At later stages (E12.5 for hChP and E14.5 for tChP), E-Cadherin signal became organized at the cell surface, displaying distinct developmental characteristics along the whole hChP and tChP (Fig. 1a). Areas adjacent to the cortical hem and rhombic lips, where progenitors of the tChP and hChP reside respectively, mostly contained cells positive for Ki-67 with only weak and diffuse E-Cadherin labeling (Pro+Dif-), indicating that local ChP epithelial cells were either still actively dividing or had just exited cell cycle^28^. Conversely, the distal ChP epithelial cells of both ventricles showed characteristics of differentiated cells, displaying only sparse Ki-67 and a strong and well-organized E-Cadherin immunostaining (Prof-Dif+) (Fig. 1a). Interestingly, Pro+Dif-areas were enriched with ChP epithelial cells positive for multiple basal bodies but mainly negative for axoneme staining. In contrast, the majority of ChP epithelial cells in Pro-Dif+ areas displayed basal body-positive structures apposed to axonemes, demonstrating that these cells were ciliated (Fig. 1b). These results illustrate that unlike ependymal cells, whose ciliogenesis starts in early postnatal ages^29^, ChP epithelial cells become multiciliated soon after they are committed (E12.5 in hChP and E14.5 in tChP)^27^. Our data show that ciliogenesis defines distinct spatial differentiation areas along the whole ChP, suggesting a temporally-regulated maturation program.

### Choroid plexus tissue expansion predominantly takes place near the ventricular wall throughout embryogenesis

To track ChP epithelial cell migration, we mapped the distribution of actively dividing cells along the ChP axis by double S-phase labeling experiments. Pregnant females were injected with both EdU and BrdU 24 hours apart, and embryos were collected 24 hours after the last injection (Fig. 2a,b). To quantify the distribution of cells that were proliferating, the ChP was divided from root to tip into four tissue segments of equal length (Fig. 2c). As expected, the percentage of EdU-positive (+) and BrdU+ cells declined over time during embryonic development. Indeed, up to 30% of the cells were proliferating at E10.5-E11.5, followed by a significant drop to less than 15% at E12.5, and a further decline to around 10% at E15.5 (Fig. 2d, e). Generally, both tChP and hChP showed a similar progression of proliferative cells along their axes. However, in contrast to data from marsupial model^25^, the mitotic nuclei were present on both dorsal and ventral surfaces of the tChP and evenly distributed from medial to lateral compartments in the hChP. Interestingly, the only stage where both ChPs showed proliferative cells distributed along all four quartiles was E12.5 (Fig. 2d, Extended data Fig. 2a, b). The percentage of proliferative cells gradually decreased along the ChP axis starting at E14.5 (Fig. 2b, d, Extended data Fig. 2b). When we inferred the proportion of dividing cells re-entering cell cycle (double positive for EdU and BrdU), we noticed that up to 40%-60% of cells possibly re-entered cell cycle over time. In addition, the peaks of cycling cells were different between the two ventricles (E12.5/E13.5 for hChP and E13.5/E14.5 for tChP) (Fig. 2f), consistent with the delayed development of tChP compared to hChP (Fig. 1). Taken together, these data suggest that growth of the tChP and hChP occurs mainly from their local roots, the number of dividing cells slows down over time and along the ChP axis, suggesting that more mature cells are usually positioned far away from the ventricular wall.

**Figure 2.**
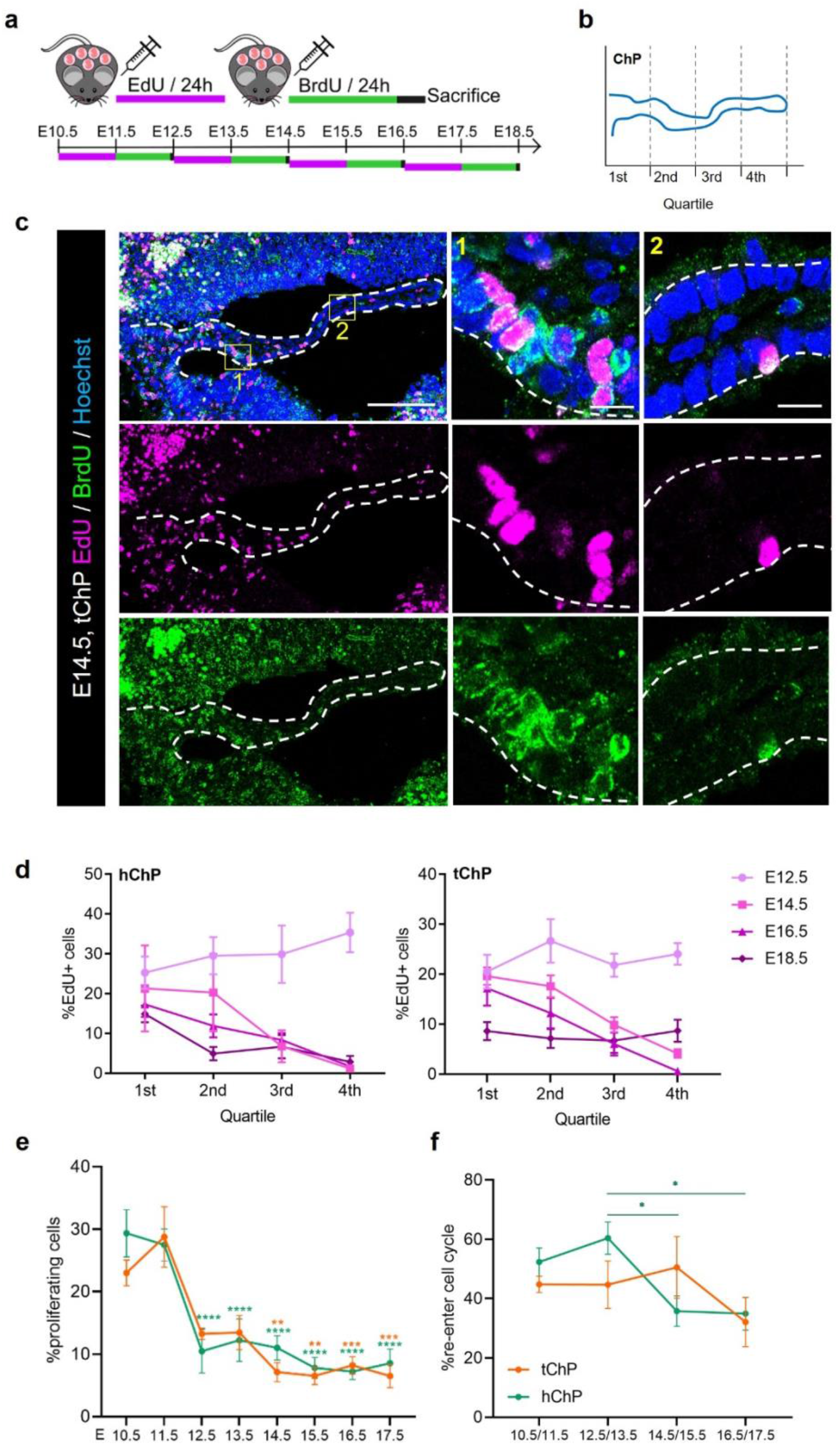
Proliferation of choroid plexus epithelial cells follows distinct spatial and temporal trajectories. **a**, Schematic representation of double S-phase protocol. **b**, E14.5 telencephalic (t) choroid plexus (ChP) sections were stained with antibodies against EdU (magenta) and BrdU (green). Insets feature ChP close to (1) or distant from (2) the ventricular wall, respectively. Scale bar: overview: 100 μm, inset: 10 μm. **c**, Schematic layout of ChP quartile partition. **d**, Quantification of percentage of EdU-positive (+) choroid plexus epithelial cells at four embryonic time points in both hindbrain (h) and t-ChP. **e**, Quantification of the percentage of total proliferative cells (EdU+ and/or BrdU+) and **f**, of the percentage of choroid plexus epithelial cells re-entering cell cycle. *n =* 3 mice. Two-Way ANOVA, *p≤ 0.05, **p≤ 0.01, ***p≤ 0.001, ****p≤ 0.0001. Bars represent mean ± SEM.

### Choroid plexus cilia are assembled in a highly organized multi-step process that matches cellular maturation

To examine the relationship between cell migration and ciliogenesis, we investigated cilia formation along the ChP length from E11.5 to late postnatal ages. ChP cilia formation was best illustrated at E16.5 in the tChP, which showed a clear gradient of differentiation along its longitudinal axis by E-Cadherin labeling (Fig. 3a). A similar gradient was also exhibited by immunostaining against AcTub, as the majority of ciliated structures were found at the tip of the tChP, far from the ventricular wall (Fig. 3b). In fact, adjacent to the ventricular wall, the majority of the ChP epithelial cells had either a single cilium (sc) or multiple cilia scattered all over the apical surface and pointed in multiple directions (ms) (Fig. 3c, d, Root and Mid). At more distal locations along the ChP longitudinal axis, most of the cells were multiciliated with cilia either scattered but oriented to the same direction (mso) or clustered and oriented (mco) (Fig. 3c, d, Mid). Cells at the ChP tip were more homogeneous and predominantly characterized by mco organization (Fig. 3c, d, Tip). These cilia configurations were confirmed by TEM ultrastructural analysis (Fig. 3e). Interestingly, cells with multiplied basal bodies but no extended axoneme, that were observed at earlier embryonic stages (Fig. 1b), were not observed at this embryonic age, suggesting that basal body multiplication happens early in the ciliogenesis process.

**Figure 3.**
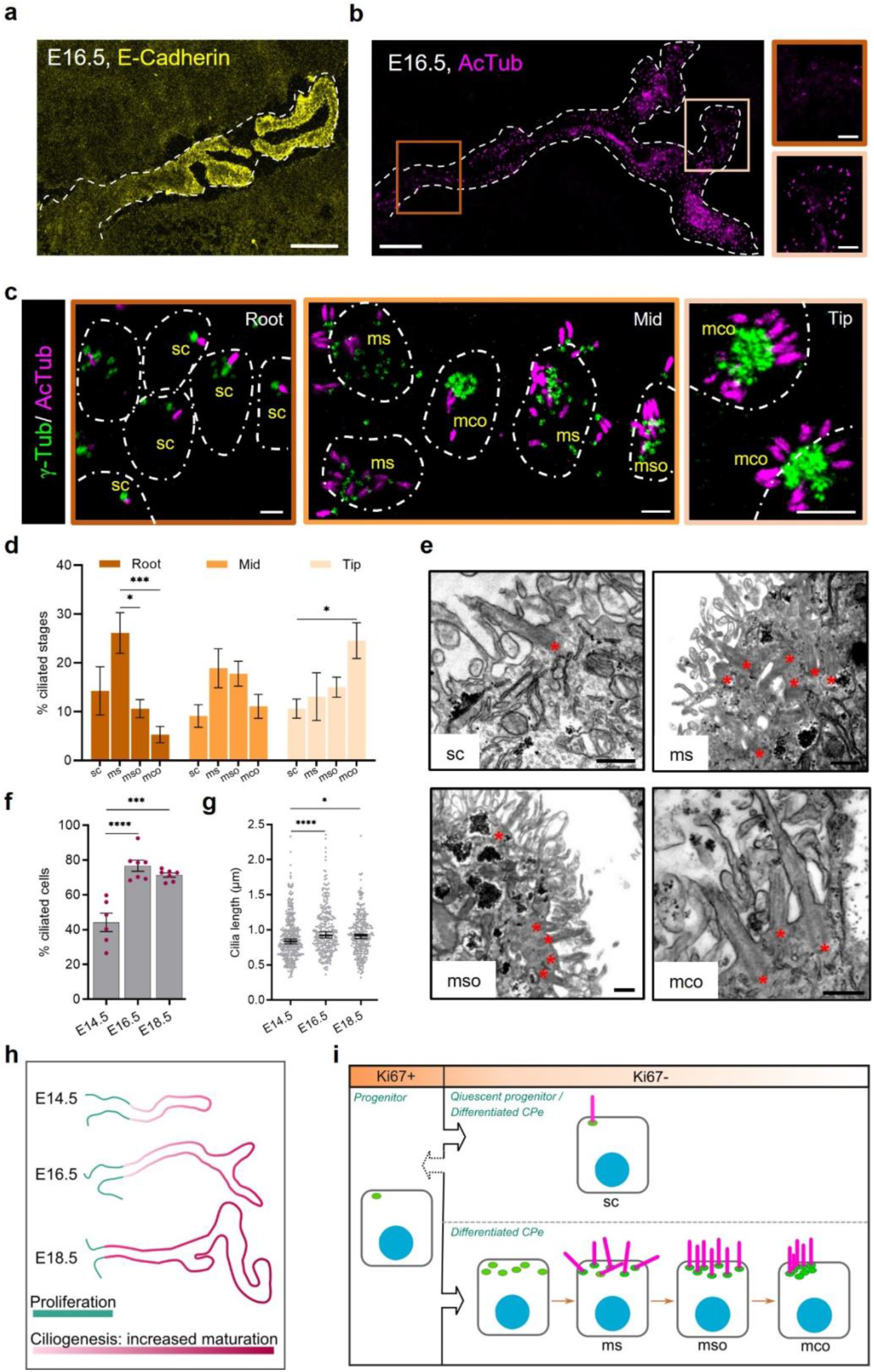
Atypical multi-step ciliogenesis in choroid plexus epithelial cells. **a, b**, Immunofluorescence images of telencephalic (t) choroid plexus (ChP) at E16.5. Sagittal brain sections were stained with antibodies against E-cadherin (yellow) (**a**) or acetylated tubulin (AcTub, magenta) (**b**). Insets feature ChP adjacent to (dark brown) or distant from (beige) the ventricular wall, respectively. **c**, ChP sections were stained with antibodies against γ-Tubulin (γ-Tub, green) and AcTub (magenta). Multiple cilia configurations were identified along the ChP according to their distance to the ventricular wall: adjacent, dark brown (Root); intermediate, light brown (Mid); distant, beige (Tip). sc: single cilium; ms: multiciliated, scattered; mso: multiciliated, scattered, oriented; mco: multiciliated, clustered, oriented. **d**, Histograms of the distribution of ciliated cells along E16.5 tChP. Bars represent mean ± SEM. *n =* 4 embryos. **e**, Transmission electron microscopy (TEM) of E16.5 choroid plexus epithelial cells showing all types of ciliated cells: sc, ms, mso and mco. **f**, Quantification of the percentage of ChP ciliated cells at three time points (E14.5, E16.5 and E18.5). Bars represent mean ± SEM. *n =* 3 mice per age. **g**, Quantification of axoneme length at E14.5, E16.5 and E18.5. Each dot represents an individual cilium. Bars represent median ± 95% CI, *n =* 250-400 cilia from 3 mice per age. **h**, Schematic overview of ciliogenesis progression in relation to spatial and temporal trajectories of cell proliferation. **i**, Schematic summary of ChP ciliogenesis. Scale bars: overview: 100 μm, inset: 20 μm (**a, b**); 2 μm (**c**); 500 nm (**e**). Two-Way ANOVA (**d, f**); Kruskal-Wallis (**g**), *p≤ 0.05, **p≤ 0.01, ***p≤ 0.001, ****p≤ 0.0001.

The presence of a single cilium in epithelial cells located at the ChP root suggested that ChP epithelial cells, as other multiciliated cells^29^, are initially monociliated. To confirm this, we stained ChP of Otx2-GFP mice against Ki67 and demonstrated that the root area primarily contained either non-ciliated proliferative Otx2+/Ki67+ cells, most likely ChP epithelial progenitors, or monociliated Otx2+/Ki67-cells (Extended data Fig. 3a, b). This suggests that ChP epithelial cells at the root possibly have the capability to re-enter the cell cycle through ciliary resorption^30^. Serial TEM reconstruction analysis of age-matched embryos confirmed the presence of monociliated ChP epithelial cells (Extended data Fig. 3c). To check whether the sc configuration was temporary, we then analyzed the cilia configuration at postnatal day (P)0. Compared to E16.5, there was a massive switch to the mco configuration. However, the percentage of sc cells was similar at the two ages, suggesting that mco is the dominant cilia configuration of mature multiciliated ChP epithelial cells, but a subset of cells remain mono-ciliated (Extended data Fig. 3d). These data indicate the presence of multiple cilia configurations in ChP epithelial cells.

During embryonic development, ChP epithelial cells tend to be initially organized as a pseudo-stratified epithelium before forming a mature monolayer^4^. We observed that when mco ChP epithelial cells were pseudo-stratified, their cilia were not consistently oriented toward the CSF. On the contrary, when mco ChP epithelial cells were arranged in a monolayer, the cilia were all pointed towards CSF (Extended data Fig. 4a-c). These data suggest an additional phase in the maturation of ChP cilia that includes a final orientation step towards the CSF compartment.

**Figure 4.**
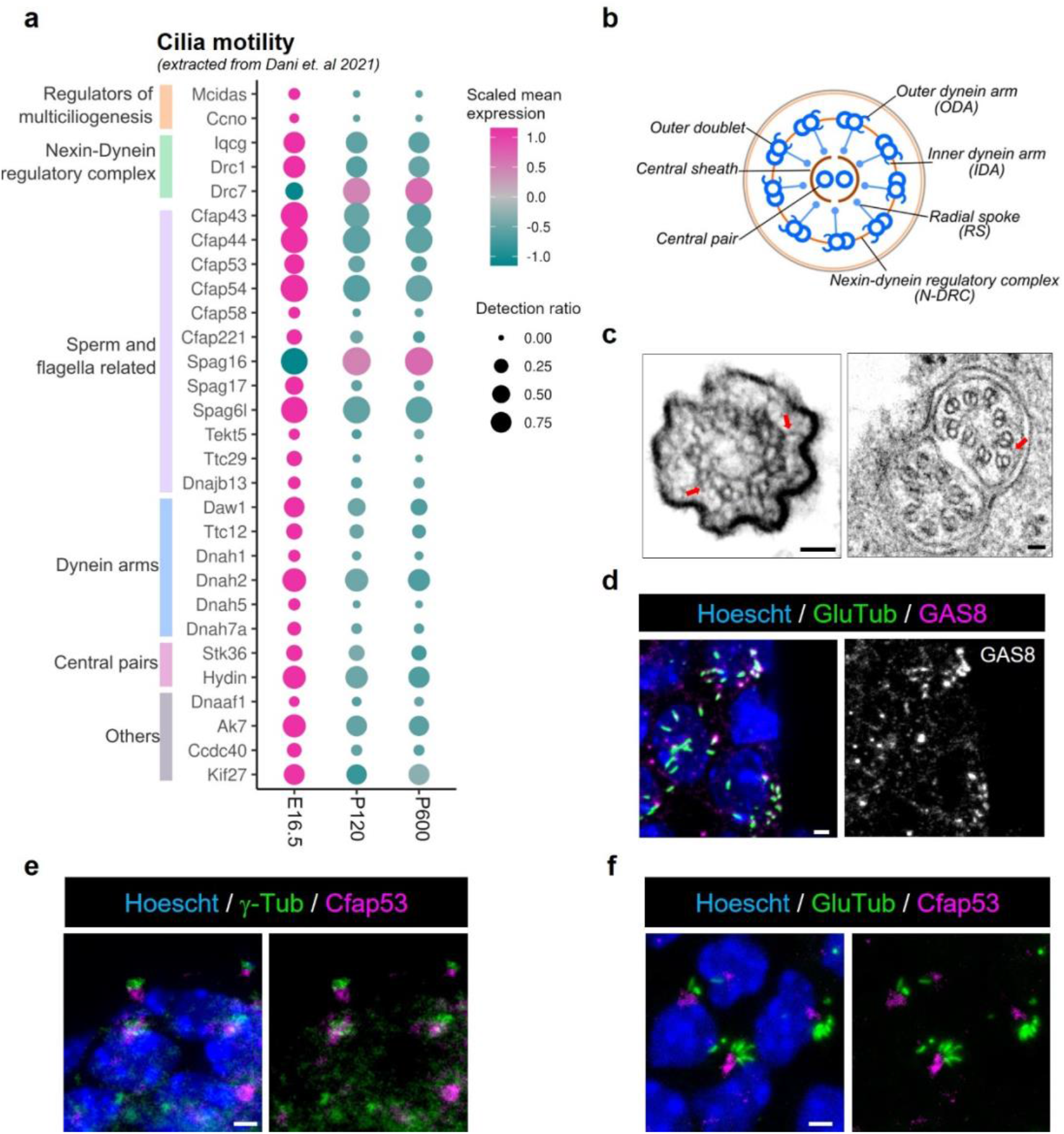
Multicilia of the choroid plexus display an atypical motile 9+0 configuration. **a**, Dot plot of cilia motility genes by snRNA-seq. Median expression level in choroid plexus epithelial cells (color) and proportion of expressing cells (circle size) of selected cilia motility candidate genes (rows). **b**, Diagram of an axonemal cross section of canonical motile cilia configuration. **c**, Transmission electron microscopy (TEM) of an axonemal cross section in choroid plexus epithelial cells at E16.5 (left) and P1 (right). **d-f**, Immunofluorescence images of E16.5 choroid plexus (ChP). ChP sections were stained with antibodies against glutamylated tubulin (GluTub, green) and GAS8 (magenta) (**d**); γ-Tubulin (γ-Tub, green) and CFAP53 (magenta) (**e**); GluTub (green) and CFAP53 (magenta) (**f**). Scale bars: 50 nm (**c**), 2 μm (**d-f**).

We then mapped the distribution of ciliated cells throughout embryonic development. At E14.5, up to 70% of ChP epithelial cells were positive for γ-Tub, showing that the cells were committed to be ciliated, however only 40 % of these cells contained a clear cilium apparatus. By the age of E16.5, up to 80% of ChP epithelial cells were ciliated. This proportion was generally maintained until birth (Fig. 3f, Extended data Fig. 4d, e). It is worthy of note that the average length of the axoneme was constant during embryonic development and ranged from around 0.9 to 1.3 μm (Fig. 3g, Extended data Fig. 4f), which is considerably shorter than ependymal (8 µm) and tracheal cilia (4-7 μm)^31^. tChP and hChP showed a similar ciliation trajectory (Extended data Fig. 4d-f).

It was previously demonstrated that tubulin post-translational modification (PTM) dynamics are closely correlated to cilia length^32^. To first evaluate the localization of the PTMs, we double-stained for AcTub and the structural microtubule marker Tubulin alpha 4a (Tub4a), demonstrating that these two antibodies were perfectly co-localized along the entire axoneme. This suggests that AcTub covers the entire length of ChP cilia (Extended data Fig. 5a). We then co-labeled for AcTub and glutamylation (GluTub, GT335), showing that all cilia contained both PTM types along the whole axoneme (Extended data Fig. 5b). These results deviate from previous description of AcTub covering longer axoneme length, including the ciliary tip, compared to GluTub, that is concentrated at the base of the axoneme^32^. Interestingly, we noticed that not all cilia were glycylated at E16.5, supporting the notion that glycylation may be mainly involved in cilia maturation rather than cilia formation in short axonemes (Extended data Fig. 5c). To note, cilia maturation in the ChP most likely is not related to axoneme length but to other unknown features^33^. Overall, our data demonstrate that ChP axonemes carry multiple PTMs and that they are post-translationally modified since the beginning of their formation.

**Figure 5.**
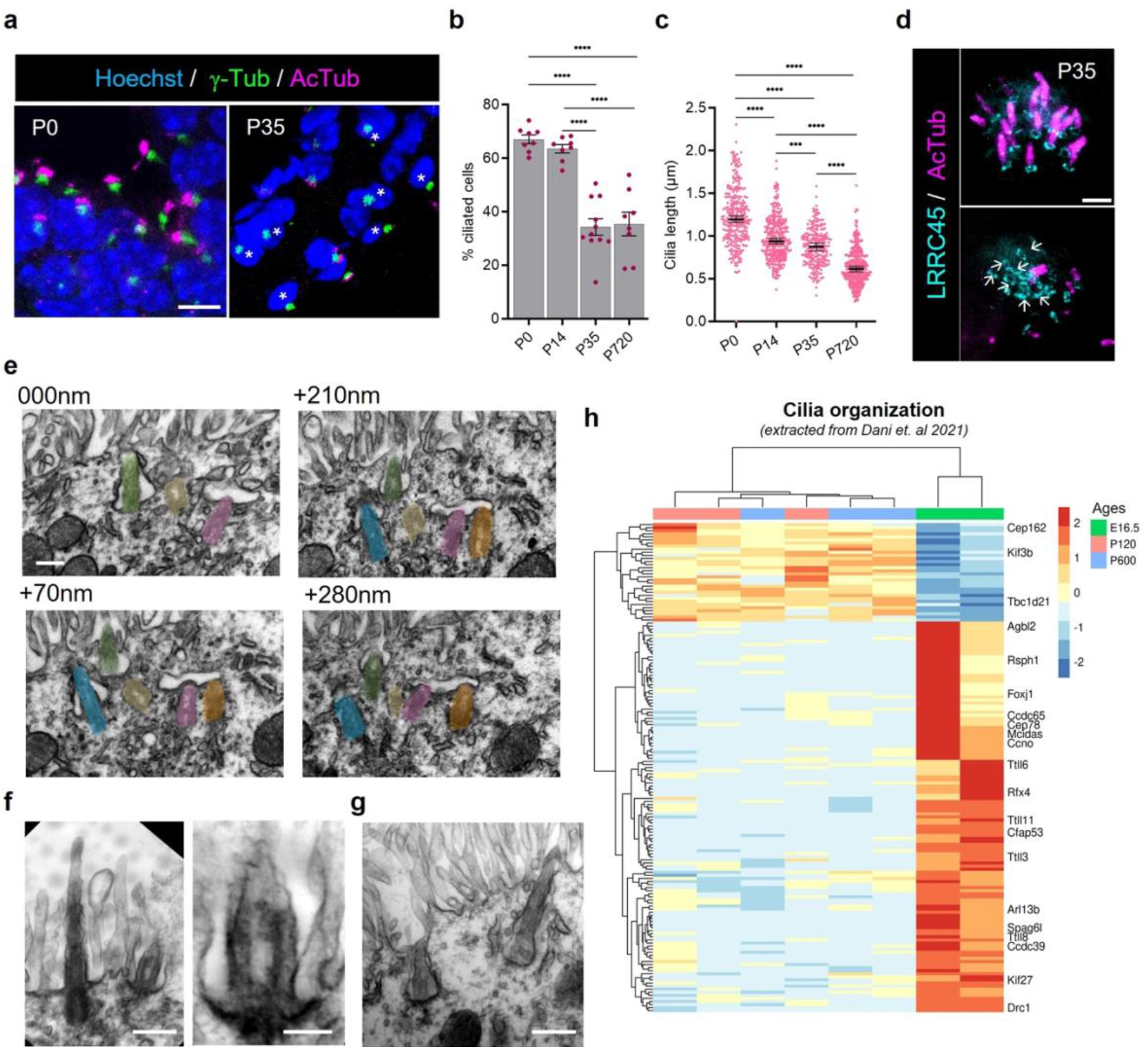
Disappearance of choroid plexus cilia starting from early postnatal ages. **a**, Immunofluorescence images of postnatal choroid plexus (ChP). ChP sections were stained with antibodies against γ-Tubulin (γ-Tub, green) and acetylated tubulin (AcTub, magenta) at P0 (left) and P35 (right). Asterisks (*) indicate choroid plexus epithelial cells negative for AcTub. **b**, Quantification of the percentage of ciliated cells at multiple postnatal ages (P0, P14, P35 and P720). Bars represent Mean ± SEM, *n =* 3 mice per age. **c**, Quantification of axoneme length at multiple postnatal time points. Each dot represents one individual cilium, *n =* 220-400 cilia from 3 mice per age. Bars represent Median ± 95% CI. **d**, Super resolution images (STED) of P35 ChP. Brain sections were stained with antibodies against AcTub (magenta) and LRRC45 (cyan). Arrows indicate single LRRC45-positive basal bodies that are negative for AcTub. **e**, Transmission electron microscopy (TEM) serial reconstruction showing the same multiciliated cell in an adult ChP. Colored regions of interests indicate individual axonemes. **f**, TEM images showing sagittal cross sections of multiple types of axonemes. **g**, TEM image showing cilia retreated deep into cytoplasm through deepening of ciliary pockets. **h**, Heatmap of cilia organization genes that are differentially expressed across ages from snRNA-seq data. Two-Way ANOVA (**b**), Kruskal-Wallis (**c**), ***p≤ 0.001, ****p≤ 0.0001. Scale bars: 10 μm (**a**); 1 μm (**d**); 200 nm (**e, f**); 500 nm (**g**).

To summarize, ciliogenesis of ChP epithelial cells occurs as a gradient of events that are highly regulated both temporally and spatially (Fig. 3h). The majority of ChP epithelial cell progenitors were located close to the ventricular wall and abundant at early stages of embryogenesis. Most of these cells were either proliferative (Ki67+) and therefore non-ciliated or quiescent (Ki67-) and monociliated (Fig. 3h, i). As cells advanced to later developmental stages towards the ChP tip, fewer proliferative cells were identified, and they became mainly multiciliated (Fig. 3h). We here suggest that multiciliation in the ChP starts with multiplication of pseudo-disorganized basal bodies and their docking along the apical surface, followed by local assembly of the axoneme, and finally orientation towards the CSF. At birth, ChP epithelial cells were mainly multiciliated with a homogeneous cilia configuration, although a minor number of cells remained monociliated (Fig. 3i).

### Choroid plexus displays atypical motile 9+0 cilia

The nature of the ChP epithelial cilia has been puzzling and not extensively studied^6^. We, therefore, took advantage of open-sourced single-nucleus RNA (snRNA) data of ChP epithelial cells at three available ages (E16.5, P120 and P600)^26^ and selectively extracted cilia-related genes (see materials and methods). As previously reported, we identified regulators of multiciliogenesis, such as *Mcidas* and *Ccno*^*6*^ and many constituents of cilia motility programs, such as components of dynein arms, central pairs, nexin-dynein regulatory complex (N-DRC) and genes of flagella structure and sperm movement (Fig. 4a). This transcriptomic profile and the fact that ChP epithelial cells are multiciliated suggest that, like ependyma, trachea and fallopian tube, ChP harbors motile cilia. Intriguingly, most of these genes were significantly downregulated at postnatal ages compared to E16.5, hinting that cilia motility may play different roles during embryonic and adult stages.

Compared to a conventional structure of motile cilia (Fig. 4b), TEM ultrastructural analysis showed a 9+0 axoneme with possible outer dynein arms (Fig. 4c), suggesting that despite having a 9+0 conformation, the ChP epithelial cilia could be motile. To resolve this identity dilemma, we used RT-qPCR to confirm the expression of key cilia motility-related genes from E16.5 ChP homogenates. Overall, all selected genes showed a decent mRNA quantity although the different genes showed different expression levels (Extended data Fig. 6a). Then, we confirmed the protein expression and localization of a subset of candidates by using immunofluorescence. We examined GAS8, a N-DRC component^34^, and coiled-coil containing protein CFAP53, an important regulator of ciliary motility^35^ (Fig. 4d-f). Most cilia were clearly positive for GAS8, which was mainly located along the axoneme shaft labeled with GluTub (Fig. 4d). Interestingly, CFAP53 was exclusively located close to the γ-Tub-positive basal bodies (Fig. 4e), as previously reported for nodal cilia^35^. Double staining of CFAP53 and GluTub (axoneme) confirmed that CFAP53 was not expressed along the axoneme, as reported for canonical 9+2 ependymal and tracheal cilia^35^ (Fig. 4f). This distribution was confirmed at later postnatal stages (Extended data Fig. 6d). Altogether, our data revealed that ChP cilia are atypical 9+0 motile cilia.

**Figure 6.**
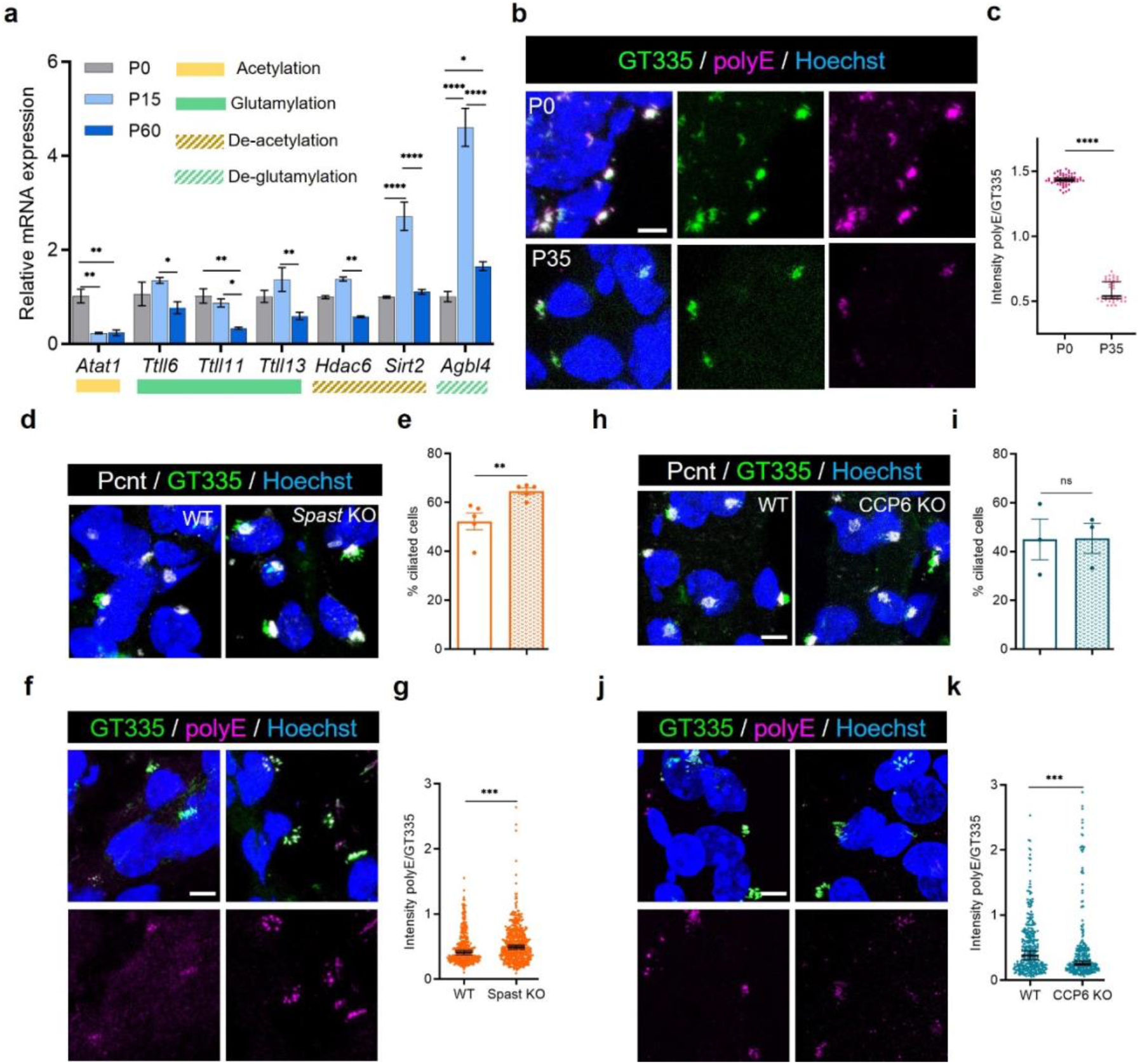
Axoneme polyglutamylation regulates choroid plexus cilia resorption through Spastin enzyme. **a**, mRNA expression level of post-translational modification catalyzing enzymes at multiple postnatal time points (P0, P15, P60). *n* = 3 mice per age. Two-Way ANOVA. **b**, Immunofluorescence images of postnatal choroid plexus (ChP). ChP sections were stained with antibodies against total-glutamylated tubulin (GT335, green) and poly-glutamylated tubulin (polyE, magenta) at P0 (top) and P35 (bottom). **c**, Quantification of poly-glutamylation level normalized to total glutamylation level at P0 and P35. Each dots represent a single cilia cluster. *n* = 40-55 cilia clusters from 3 mice per age, Mann-Whitney test. **d**, Adult *Spastin* (*Spast*) KO ChP sections were stained with antibodies against GT335 and pericentrin (Pcnt, grey). **e**, Quantification of the percentage of ciliated cells in *Spast* KO compared to control littermates (WT). *n* = *5* mice per genotype, Mann-Whitney test. **f**, Adult *Spast* WT and KO ChP sections were stained with antibodies against GT335 and polyE (magenta). **g**, Quantification of poly-glutamylation level normalized to total glutamylation level in *Spast* KO compared to littermate WT. *n =* 250-350 cilia cluster from *5* mice per genotype, Mann-Whitney test. **h**, Adult *Agbl4* knockout (CCP6 KO) ChP sections were stained with antibodies against GT335 and Pcnt. **i**, Quantification of the percentage of ciliated cells in CCP6 KO compared to WT. *n* = *3* mice per genotype, Mann-Whitney test. **j**, Adult CCP6 WT and KO ChP sections were stained with antibodies against GT335 and polyE. **k**, Quantification of poly-glutamylation level normalized to total glutamylation level in CCP6 KO compared to WT. *n =* 280-320 cilia cluster from 3 mice, Mann-Whitney test. Bars represent Mean ± SEM (**a, e, i**); Bars represent Median ± 95% CI (**c, g, k**). Scale bars: 5 μm. *p≤ 0.05, **p≤ 0.01, ***p≤ 0.001, ****p≤ 0.0001. ns: not significant.

### Gradual postnatal disappearance of choroid plexus cilia

ChP epithelial cells go through critical rearrangements in the postnatal period^36,37,38^. We, therefore, decided to map ChP cilia remodeling at birth (P0), P14 (eye opening), P35 (end of synaptogenesis) and P720 (aged brain). Quantification of the percentages of cells containing basal body clusters using γ-Tub showed that the majority of ChP epithelial cells were positive at birth (up to 95%). Starting from P35, the percentage of positive cells mildly decreased and then remained plateaued until P720 (around 70%) (Extended data Fig. 7a). Surprisingly, the percentage of ciliated ChP epithelial cells drastically decreased starting from P35 (almost 50%). No further decrease was observed in aged P720 ChP (Fig. 5a, b). Both ventricles followed a similar trend, suggesting that the loss of axonemes is a systemic event for the ChP (Extended data Fig. 7b). We then measured the length of the axonemes, noticing that length peaked at birth (1.2 ± 0.02 µm) and then steadily decreased, reaching the lowest point at P720 (0.6 ± 0.01 µm) (Fig. 5c, Extended data Fig. 7c). These results showed that ChP epithelial cilia are dynamic apparatuses that may adapt to different circumstances through the mouse life. To verify the axoneme disappearance, we applied both super-resolution (STED) and ultrastructural (TEM) microscopic analysis. Using LRRC45, a marker for the distal end of the mother centriole^39^, to label individual basal bodies, we identified two major types of cilia clusters: one where all basal bodies contain clear axonemes (Fig. 5d, top) and one where most of the basal bodies are not apposed to axonemes or display only stubby, truncated axonemes (Fig. 5d, bottom). TEM serial analysis revealed a more complex situation. Within the analyzed ciliated cells (*n* = 613 total cells, 3 mice), only 8% of the total measured axonemes (*n* = 205 total axonemes, 3 mice) showed a complete structural configuration characterized by a large basal body, thinner axonemal tip and clear ciliary tip^40,41^. The remaining analyzed basal bodies had at least one axoneme (97.6%, *n* = 613 total cells, 3 mice), which however looked exceptionally short and with an extended ciliary tip (Fig 5e, f; Extended data Fig. 8a-c). These observations leave an open question on how exactly tubulin is organized in these cells. We also observed that frequently cilia from adult ChP were deeply embedded in the cytoplasm and surrounded by defined ciliary pocket (CiPo) or that the stubby axonemes were surrounded by a structure resembling an intracellular vacuole, probably by enclosure of CiPo (Fig. 5g, Extended data Fig. 8b). CiPo is a clathrin-coated pit closely connected with the actin-based cytoskeleton and is a dynamic center for endocytosis^42^. It has been demonstrated that resorption of primary cilia depends on CiPo and endocytosis^43,44^, suggesting that the dismantling of multiple cilia in ChP may involve endocytosis of individual cilia through ciliary pockets^45^.

**Figure 7.**
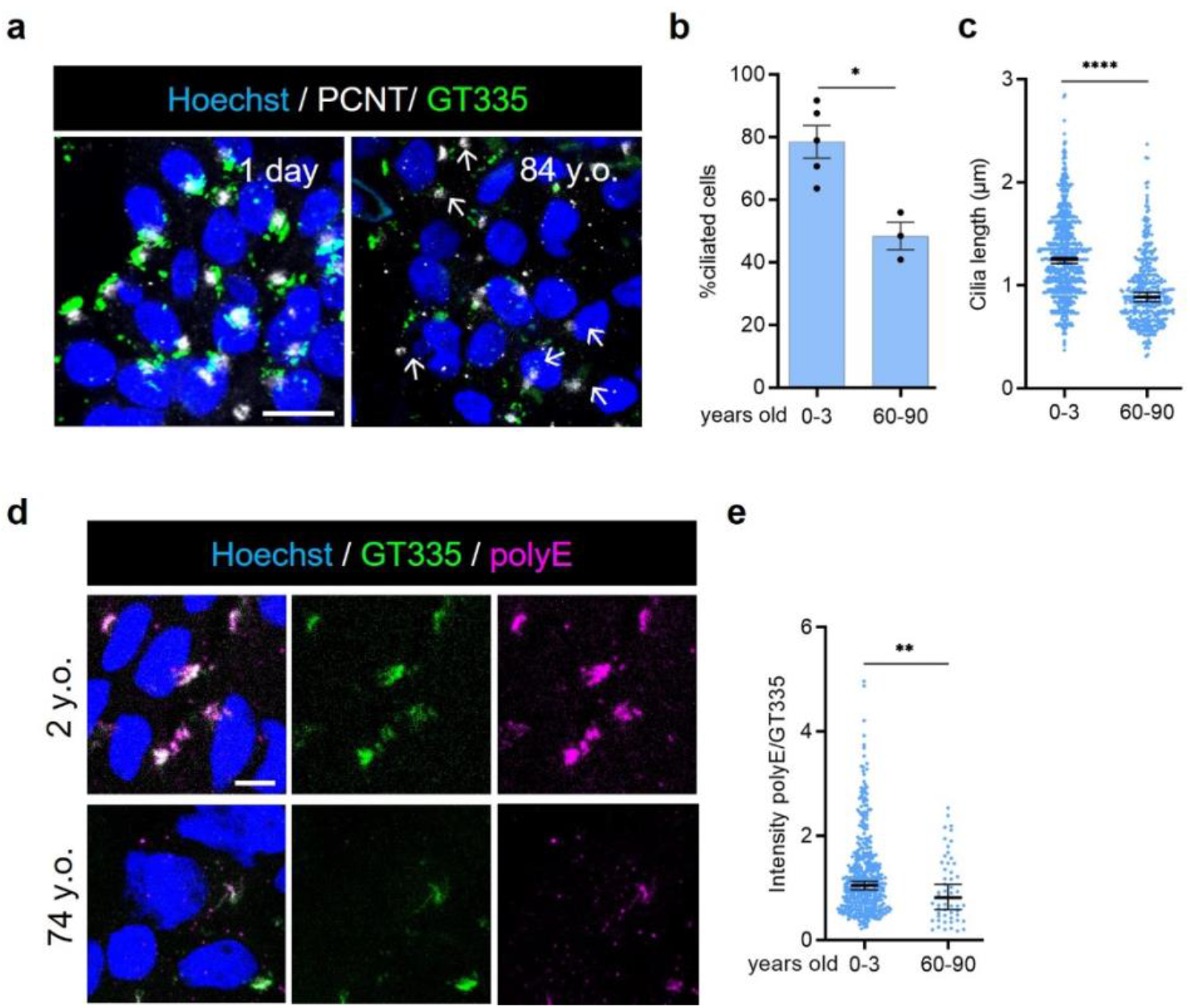
Resorption of human choroid plexus cilia is regulated by polyglutamylation levels. **a**, Immunofluorescence images of human postmortem choroid plexus (ChP). Representative ChP sections stained with antibodies against glutamylated tubulin (GT335, green) and pericentrin (PCNT, grey) in 1-day-old patient (left) and in 84-year-old patient (right). **b**, Quantification of the percentage of ciliated cells in young patients (0-3 years-old) and aged patients (60-90-year-old). Bars represent Mean ± SEM, *n =* 5 young specimens, 3 aged specimens, Mann-Whitney test. **c**, Quantification of axoneme length in young and aged patients. Bars represent Median ± 95% CI. Each dot represents one individual cilium, *n =* 420-800 cilia from 3-4 patients per age group, Mann-Whitney test. **d**, Representative ChP sections were stained with antibodies against GT335 (green) and poly-glutamylated tubulin (polyE, magenta) in 2 years-old patient (top) and in 74-year-old patient (bottom). **e**, Quantification of poly-glutamylation level normalized to total glutamylation level in young and aged samples. Each dots represent a single multicilia complex. *n =* 3 samples per age. Bars represent Median ± 95% CI. Mann-Whitney test. Scale bars: 10 μm (**a**), 5 μm (**d**). *p≤ 0.05, **p≤ 0.01, ****p≤ 0.0001.

At the transcriptional level, the snRNA data from Dani et. al^26^ showed that the vast majority of cilia genes (curated list based on GO Cilia organization) had reduced expression at P120 and P600 compared to E16.5 (Fig. 5h) - a trend that was maintained across ventricles (Extended data Fig. 7d). Altogether, we demonstrated that ChP epithelial multicilia are not maintained throughout life but rather disappear at an early postnatal stage, which is quite a unique developmental characteristic of ciliated cells.

### Destabilization of microtubules is upstream of choroid plexus cilia resorption

While cilium assembly is a well characterized process, much less is known about the mechanism of cilium disassembly/resorption and most of the available knowledge is based on studies of primary cilia^45^. We hypothesized that microtubule destabilization could represent an important step in the resorption of the axoneme. Multiple PTMs have been linked to microtubule turnover and tubulin destabilization^45, 46,47^. We, therefore, examined the mRNA expression levels of key enzymes regulating PTMs throughout ChP development and noticed quite a unique pattern of expression (Extended data Fig. 9a, b). As expected, the majority of the enzymes were highly expressed embryonically and decreased at birth. When focusing on postnatal periods, we observed a significant down-regulation of the acetylase *Atat1* at P15 compared to P0. Interestingly, *Atat1* expression levels remained low afterwards, indicating diminished tubulin acetylation in adult animals (Fig. 6a). On the contrary, glutamylases and glycylases, members of the *Ttll* genes, tended to be constantly expressed from birth to P15 and then significantly decreased at P60. *Ttll6*, -*11*, -*13* polyglutamylases showed the strongest differences (Fig. 6a). Surprisingly, enzymes relating to cilia disassembly, such as deacetylases *Hdac6, Sirt2* and deglutamylase *Agbl4* (encodes for CCP6) were all selectively upregulated at P15 compared to P0, and their expression levels returned to basal level at P60. These results strongly suggest the existence of a specific developmental time window when ChP epithelial cells activate a transcriptional pathway dedicated to cilia resorption, which will manifest at P35 (Fig. 5a, b). We, then, used selective antibodies to measure the intensity level of the total glutamylation (GT335)^48^ and polyglutamylation (polyE, side chains longer than four glutamates)^49^. Remarkably, the ratio between polyglutamylation/total-glutamylation (polyE/GT335) significantly decreased at P35 compared to P0, suggesting a drastic destabilization of tubulin at this stage (Fig. 6b).

To understand whether the level of tubulin glutamylation is essential for axoneme maintenance in ChP epithelial cells, we first focused on spastin, a microtubule-severing protein that breaks longer microtubules into shorter ones^21,22,50^. Spastin severing activity is a biphasic event that is initially activated by an increase in tubulin polyglutamylation. However, further accumulation of glutamates inhibits spastin activity^22^. To test the involvement of spastin in ChP epithelial cilia stability, we analyzed P35 spastin KO (*Spast* KO) mice. Interestingly, these mice were characterized by a subtle but significant increase of the percentage of ChP epithelial ciliated cells and by a significant increase of the poly-glutamylation ratio compared to wild-type (WT) littermates (Fig. 6d-g), suggesting steadier axoneme microtubules. To note, *spast* mRNA did not change throughout development (Extended data Fig. 9c), indicating that our phenotype is mainly regulated at local protein level. We, therefore, hypothesized that increase of glutamylation level may be sufficient to control axoneme stability. To test this idea, we considered CCP enzymes, as they remove glutamate residues from the C-terminal tails controlling the length of glutamylate chains on tubulin^49^. Having identified *Agbl4* as the enzyme with the highest expression in the ChP at P15 (Fig. 6a; Extended data Fig. 9b), we analyzed *Agbl4* knockout (KO) mouse (CCP6 KO)^51^. CCP6 KO mice did not show major differences when compared to WT littermates, as the percentage of ciliated cells was unchanged and the poly-glutamylation ratio was still reduced in the CCP6 KO (Fig. 6h-k). These data demonstrated that CCP6 is not essential either for cilia formation or cilia maintenance in the ChP, however it could be that other CCPs may have compensated for the loss of this single enzyme. In conclusion, we demonstrated that the level of tubulin polyglutamylation is key for maintaining ChP cilia and that lack of spastin generated a delay of cilia resorption.

### Cross-species ciliary tubulin phenotypes in postmortem human choroid plexus

Finally, to expand the relevance of our data, we processed human postmortem ChP specimens at multiple time points. In young ChP (Extended Table 1) up to 80% of ChP epithelial cells, identified as positive for OTX2, TTR and E-Cadherin (Extended data Fig. 10a), were ciliated, as demonstrated by double staining for pericentrin (PCTN) and GT335. This percentage significantly decreased to 60% in aged ChP specimens (Extended Table 1) (Fig. 7a, b, Extended data Fig. 10b). Similarly, the total axoneme length significantly decreased over time (Fig. 7c). These results demonstrate that the disappearance of cilia, together with reduced axoneme length, are conserved between rodent and human ChPs. Interestingly, the level of tubulin polyglutamylation significantly decreased in aged human ChP (Fig. 7d, e), demonstrating that a similar molecular apparatus is involved in the control of tubulin destabilization in human and mouse.

## Discussion

In this manuscript, we have characterized the biogenesis and maintenance of multiciliated epithelial cells in murine ChP. Our work showed that embryonic ChP proliferative cells progress on both sides of the growing plexus and that ChP epithelial ciliation states are closely correlated with cellular proliferation and differentiation stages. Uniquely, the majority of the ChP axonemes are resorpted during the postnatal period, most likely through an intrinsic destabilization of the microtubular cytoskeleton driven by downregulation of polyglutamylation. This effect is mediated in part by the microtubule-severing protein spastin. Finally, we showed that axoneme resorption is not a species-specific event, as it also happens in postmortem human ChP epithelial cells.

Cilia formation and maintenance are well-known processes historically studied in unicellular models and/or *in vitro* cell cultures^30^. We here studied ciliogenesis in the murine ChP *in vivo*, confirming that epithelial cells of the ChP are multiciliated and that cilium formation is entirely developed during embryogenesis. Based on the lack of deuterosome-like structures in our E16.5 TEM reconstructions and the recent knowledge that deuterosomes can be dispensable from multiciliogenesis^52,53^, we hypothesize that ChP epithelial ciliogenesis is a parental centriole-dependent pathway that would allow the production of up to 10-20 centrioles^54^. We also show that the number and length of ChP epithelial axonemes fluctuate over time, peaking at birth, which suggests that the role of the cilia may change throughout development.

Detailed transcriptomic analysis demonstrates that the ChP expresses high levels of well-known motility cilia genes at E16.5 and that their expression decreases postnatally. Interestingly, ChP cilia contain key N-DRC components, like GAS8 and CFAP53 - a protein known to be expressed only in motile cilia, where it links outer dynein arms to other molecular axoneme components^35^. Although we confirmed that the murine ChP axonemes have a 9+0 configuration from embryonic to adult mice, we noticed that CFAP53 is selectively located close to basal bodies of the ChP cilia, suggesting the possibility of ciliary rotational movements^35^, if any. Another interesting phenotype that supported possible motile functions of the cilia was the incomplete orientation of their axonemes during embryonic stages, as already reported for mucociliary clearance cells. In this cell type, the final axonemal reorientation is directly correlated to the presence of the dynein arms and dependent from the flow direction^55,56^. It is, however, important to notice that functional studies related to the ChP cilia have suggested that these cilia may act as primary cilia. For example, the analysis of one-week-old ChP of *Tg737*^*orpk*^ mutants showed elevated intracellular cAMP levels and altered pH regulation and ion transporter activity^57^. It is therefore possible that ChP cilia may temporally act as primary cilia after birth, but it remains possible that their functions and properties may change over time or that ChP cilia are heterogeneous. Due to the lack of a valid *in vitro* ChP model that will recapitulate all ciliogenesis stages, the analysis of some of these critical steps can become quite difficult and limited by temporal resolution. However, in the future, it will be important to evaluate ciliogenesis progress in a cell-by-cell model.

One of the major phenotypes that we identified in our study is the atypical progressive resorption of cilia in both mouse and human ChP. To note, not all cilia disappeared from the ChP, suggesting that local cilia may have different functions at different developmental stages. We demonstrated that the resorption was induced by local microtubule dismantling coordinated by the severing-enzyme protein spastin. The loss of *spastin* expression in *Drosophila* or the hyperactivity of *spastin* in *in vitro* models is known to disrupt axonal microtubule stability^58,59,60^. Further, spastin-dependent microtubule destabilization is essential for physiological pruning of motor axon branches from the neuromuscular junctions^50^. As far as we know, our study demonstrated, for the first time, that spastin can also regulate cilia assembly and disassembly *in vivo* in the ChP. These data are essential taking into consideration that mutations in the *SPAST* gene are responsible of causing hereditary spastic paraplegia (HSP). Like multiple ciliopathies, HSP patients can be characterized by cognitive deficits and declined intellectual capabilities^15,61,62^, suggesting that in the future it will be important to expand our spatio-temporal understanding of tubulin-associated genes in non-neuronal cells. Intriguingly, dementia and reduced intelligence quotient are late-onset symptoms arising from 40-50 years of age onward and data from our lab (not shown) and from postmortem human ChP^38^ proved that the ChP goes through critical morpho-functional reorganization starting from middle age stages.

Polyglutamylation is essential for ciliary maturation^63^ and proper beating in motile cilia^64^. Our data, for the first time, demonstrate that the level of polyglutamylation may also control the very existence of a fully developed axoneme in a *in vivo* model, most likely generating or modulating local signals of selective target proteins, like the recruitment of spastin^21,22^. Our data showed that *spast* mRNA level is stable over time, indicating that axoneme resorption depends on local spastin protein whose activity is modulated by the length of the tubulin glutamate-side chain. In fact, deletion of *spast* increased the percentage of ciliated cells by restoring tubulin polyglutamylation level. Our early transcriptomic data showed that already at the age of P15, nearly 3 weeks before observing cilia disappearance, there is a significant increase in the expression of *Agbl4* deglutamylase, involved in removing tubulin PTMs, suggesting that physical axoneme resorption requires a decent amount of time to be completed. While the binding affinity of spastin to microtubules linearly increases with the level of glutamylation, spastin enzymatic activity only increases when the number of glutamates rises from one to eight and decreases when more glutamate residues are added^22^. We, therefore, reason that at embryonic stages and at birth, high level of polyglutamylation may inhibit the severing activity of spastin, despite the fact that the enzyme has high affinity to tubulin. At P15, accelerated levels of *Agbl4* may remove enough glutamates from the tubulin chain, allowing spastin to reach its full severing capacity, destabilizing the axonemes. Another possibility could be the presence of unknown cofactors that may independently activate and/or control spastin activity. Little is known about the molecular apparatus related to spastin that is involved in microtubule dynamics. One of the main known groups of proteins are the collapsing response mediator proteins (CRMP1-5), that have been shown to bind spastin and enhance its function in neuronal cells^65^. Thus, it is important to understand the cellular mechanism of microtubule severing enzymes and how their activities are spatially and temporally restricted. Curiously, we show that the expression level of TTLL enzymes were stable until adult age and then significantly downregulated, suggesting a possible general reduction of the cilia polyglutamylation process. Deficit in TTLL expression level does not affect axoneme structure but has been correlated to severe defects of sperm motility and shortening of the sperm distal end^66^.

Altogether, our study identifies polyglutamylation as a direct cause of axoneme resorption in the ChP, bolstering the importance of tubulin PTMs in brain physiology. While the aberration of tubulin PTMs, as polyglutamylation and acetylation, is sufficient to induce neurodegeneration^32,67,68,69^, no unifying molecular mechanism underlying microtubule stability has so far been found. Future experiments will be essential to uncover the specific role of tubulin in the ChP^13^ and how the fate of ChP ciliated cells is regulated, in particular the mechanisms involved in cilia function and maintenance.

## Materials and Methods

### Choroid plexus human samples

De-identified ChP specimens were obtained from the NIH Neurobiobank at the University of Maryland, Baltimore, MD. All tissues and experiments were handled in strict accordance with ethical guidelines and regulations for the research use of human tissue set forth by the Medical Faculty of the Heidelberg University (S-753/2019). Age at the time of tissue collection is stated in Supplementary Table 1.

### Experimental animals

Inbred C57BL/6J and C57BL/6N mice were purchased from Janvier labs. GFP-tagged *Otx2* mouse line was kindly provided by Dr. Thomas Lamonerie^70^. Animal were housed in both the Interdisciplinary Neurobehavioral Core (INBC) at Heidelberg University and the German Cancer Research Center (DKFZ) animal facility. Both male and female mice were included in the study. Animals were housed in static cages, no environmental enrichment toys were added to the cages, only extra Kimwipes tissue. After weaning, animals were socially housed in groups of two to five per cage. Animals were housed in a temperature and humidity-controlled room on a 12 hours light/12 hours dark cycle and had food and water *ad libitum*. All animal experiments were approved by the local governing regional council (*Regierungspräsidium Karlsruhe*, Germany). All methods were carried out following the German Animal Welfare Act regulations. Animal studies are reported in compliance with the ARRIVE guidelines.

P35 *Spastin* knockout (KO) mice were generated as described by Brill et al. 2016^50^ and housed at the Institute of Neuronal Cell Biology, Technische Universität München, Munich, Germany. Experiments on *Spastin* KO animals were performed in accordance with the regulation of the Government of Upper Bavaria. P60 *Agbl4* (CCP6) KO mice were generated as described by Magiera et al. 2018^51^ and housed at the Institut Curie, Orsay, France. Experiments on *Agbl4* KO animals were approved by the ethics committee of the Institut Curie CEEA-IC #118 (authorization no. 04395.03 given by National Authority) and performed in accordance with the recommendations of the European Community (2010/63/UE).

### Double S-phase labeling

Time mated females were checked for the presence of a vaginal plug every morning between 6:30 and 7:30 AM. E0.5 was determined as midday of the day when vaginal plug was detected. To confirm pregnancy, body weight of females was collected every other day. E10.5-E16.5 pregnant dams received intraperitoneal (i.p.) injection of EdU (#A10044, Invitrogen) at around noon, followed by a second injections 24 hours later BrdU (#B5002, Sigma). Embryos were collected 24 hours after BrdU administration. EdU and BrdU were freshly prepared on injection days by dissolving in 50°C sterile PBS to 6.15 mg/ml^71^. The dosage for each Uridines was 100 mg/kg body weight.

### Tissue processing for confocal and STED microscopies

Whole embryos (E11.5-E12.5), whole heads (E13.5 – E16.5) and dissected brains (E18.5 – P720) were rinsed in cold PBS, embedded in Tissue-Tek OCT and then frozen on dry ice. Double S-phase labeling tissues were first rinsed in cold PBS, immersed in 4% PFA for 2 hours, cryoprotected overnight in 30% sucrose and finally embedded in OCT and frozen on dry ice.

#### Immunostaining

All tissues were sliced in 15 µm-thick coronal or sagittal sections with a cryostat (Leica CM1950). Before immunostaining, frozen sections were fixed with cold 4% PFA on ice, blocked and permeabilized (10% donkey serum and 0.5% Triton X-100 in PBS) for 1 hour, then incubated in primary antibodies overnight at 4°C followed by secondary antibodies for 2 hours (3% donkey serum and 0.5% Triton X-100 in PBS). Sections were then counterstained with Hoechst 33342 (Biomol, #Cay15547-25) and mounted with Dako mounting medium (#S302380-2, Agilent). The following primary antibodies were used: rabbit anti-Ki67 (1:700, #PA5-19462, Thermo Fisher), mouse anti-E-Cadherin (1:200, #610182, Becton Dickinson), mouse anti-γ-tubulin (1:1500, #MA1-19421, Invitrogen), rabbit anti-acetylated α-tubulin (1:800, #5335T, Cell Signaling), mouse anti-glutamylated tubulin GT335 (1:800, AdipoGen #AG-20B-0020), rabbit anti-pericentrin (1:200, #ab4448, Abcam), rabbit anti-GAS8 (1:200, #HPA041311, Atlas Antibodies), rabbit anti-RSPH9 (1:200, #HPA031703, Atlas Antibodies), rabbit anti-CFAP53 (1:200, #A16607, ABclonal), rabbit anti-glycylated tubulin (1:5000, #AG-25B-0034-C100, Biomol), chicken anti-GFP (1:1000, #GFP-1010, Hölzel), rabbit anti-polyE (1:1000, #AG-25B-0030-C050, AdipoGen), mouse anti-α-tubulin (1:1000, #T9026, Sigma-Alrich), guinea pig anti-LRRC45 (1:200, kindly provided by Dr. Gislene Pereira), Secondary antibodies were selected from Alexa Fluor 488/555/594/647 donkey (1:500, Jackson ImmunoResearch). For Stimulated Emission Depletion Microscopy (STED) microscopy a goat anti-guinea pig secondary antibody coupled to Abberior STAR 635 (1:500, Abberior GmbH) was used.

#### Double S-phase labeling

PFA-fixed sections were fixed a second time with 4% PFA for 10 minutes and then permeabilized with 1% Triton X-100 for 1 hour. EdU detection was performed using Click-iT EdU Imaging Kit (#C10339, Invitrogen) following manufacturer’s protocol. Tissue was, then, incubated with 37°C 2N HCl for 40 minutes to denature DNA for BrdU labeling. HCl was neutralized with 1M Tris pH8.0. Afterwards, tissue was blocked and permeabilized (see above) and incubated with anti-BrdU (1:500, #BU1/75, Abcam) overnight at 4°C. For secondary antibody labeling and later steps see above.

### Confocal and super-resolution microscopy acquisition and analysis

Confocal microscopy images were acquired using a Zeiss LSM780 microscope with x20, x40 or x63 objective (oil immersion). Images were acquired with 2048 × 2048-pixel. ZEN Black software was used for image acquisition. Images used for polyE quantification were acquired with a Leica SP5 microscope with x63 objective. Images were acquired with 2048 × 2048 pixel.

#### Double S-phase

whole ChP tissue was acquired using 40x oil objective with tile-stitch module of 10% overlap and z-stack projection of 3-5 μm range. From the ventricular wall to the tip of the ChP tissue, 4 sections of equal length were defined for quantify the total number of ChPe, calculated as EdU+ and BrdU+ cells. The percentage of cells re-entering cell cycle was calculated as (EdU+ and BrdU+)/EdU+ cells.

#### Cilia apparatus

Selective ChP regions (bottom, mid and tip based on distance from ventricular wall) were acquired with 63x oil objective, zoom 2, z-stack projection of 0.35 μm interval. All images were acquired for the entire penetration of the antibodies to ensure the full reconstruction of cilia apparatus. Cell density, number of axonemes and basal bodies were quantified using Cell Counter plug-in. Cilia length was manually measured in max intensity projection images. For GT355 and polyE expression analysis the mean pixel intensity of the single channel signal in each field was measured using Fiji (https://imagej.net/)^72^.

Super-resolution/STED microscopy images were acquired with an Abberior Instruments FACILITY LINE STED microscope equipped with an inverted IX83 microscope (Olympus), a 60x oil objective (UPlanXApo 60x/1.42 oil, Olympus), using pulsed excitation lasers at 561 nm and 640 nm and a pulsed STED laser operating at 775 nm at 40-MHz repetition rate. All acquisition parameters were controlled via the Lightbox Software (Abberior Instruments). All images were processed and analyzed with ImageJ (Fiji, NIH)^72^.

### Transmission electron microscopy (TEM)

E16.5 (*n* = 4) and P60 (*n* = 3) animals were fixed with 2.5% glutaraldehyde in cacodylate buffer (CaCodylate buffer 50mM, KCl 1M, CaCl_2_ 0.1M, MgCl_2_ 0.1M, 2.5% Glutaraldehyde and 2% sucrose). The ChP was dissected and post-fixed for 1-2 hours in the same fixative at 4°C. After washing in PB 0.2 M, the ChP was post-fixed in osmium ferrocyanide (1 volume of 2% aqueous osmium tetroxide: 1 volume of 3% potassium ferrocyanide) for 1 hour at 4°C, dehydrated for 15 minutes in increasing concentrations of acetone (30%, 60%, 90%, 100%), incubated in acetone/Spurr resin (1:1, 30 minutes, 1:2, 30 minutes) and in Spurr resin (Sigma-Aldrich) at room temperature (RT) overnight. Finally, the ChP was embedded in Spurr resin in capped 00 BEEM capsules (Electron Microscopy Sciences) for 24 hours at 70°C. Ultrathin sections were cut with an ultramicrotome (EM UC6, Leica Microsystems), collected on uncoated nickel grids (100 mesh) and counterstained for 30 seconds with UranyLess EM Stain and 30 seconds with lead citrate (Electron Microscopy Sciences). Sections were observed with a JEM-1400 Flash transmission electron microscope (JEOL, Tokyo, Japan), and images were acquired with a high-sensitivity sCMOS camera.

Epithelial choroid plexus cells (n = 613 in total; 3 P60 mice) were selected such that both apical and basal borders of the cells were visible, and nucleus was at its best fit. The cilia quantification analysis was performed manually with ImageJ (Fiji, NIH)^72^.

### qRT-PCR

RT-qPCR using SYBR was performed with the whole procedure from tissue isolation to data analysis as described in detail previously^73^. Briefly, tChP and hChP tissue was isolated and pooled together for RNA extraction. 3 biological replicates were included in each age. Due to the small size of E16.5 tissue, samples from 3 embryos were pooled for each biological replicate. Non-baseline-corrected raw data was processed with LinRegPCR (version 2020.0)^74^ to obtain amplification efficiency (E) and quantification cycle (Cq) for relative quantification. The software also returned starting concentrations (N0, arbitrary fluorescence unit) for absolute quantification in Fig. 3. *Hprt, Rpl27* and *Rpl13a* were used as reference genes. Detailed primer sequences are in Supplementary Table 2.

### snRNA sequencing analyses

For snRNA sequencing data analysis, the *matrix, features* and *barcodes* files were downloaded from NCBI GEO (GSE168704)^26^ and reconstructed into a SingleCellExperiment object using *DropletUtils. SingleCellExperiment* package was used for storing and retrieving data from the S4 object. Data quality control, dimensionality reduction and visualization was performed using package *scater*. Data normalization and modeling gene variance by expression was performed with *scran*, the top 2000 most variable genes were selected to perform Principal Component Analysis (PCA). As suggested in the original paper, X inactivation and Y chromosome genes were removed. The top 10 PCA were used for UMAP dimension reduction with n-neighbors = 15. Cell annotations were retained from the original analysis in Cell Metadata. All epithelial cells were subtracted and used for subsequent analyses. *DESeq2* package with shrinkage method lfcShrink type = ashr^75^ was applied to analyze differentially expressed genes (DEG) of epithelial cells. The gene list of GO term Cilium Organization (GO: 0044782) was expanded by adding selected genes, including cilia PTMs and cilia disassembly (p-value < 0.05). The same DEG analysis pipeline was performed for 3 ventricles, collectively and separately.

### Statistical analysis

Statistical analysis was performed using GraphPad Prism (version 9.3.1). Appropriate statistical tests were selected based on the distribution of data and sample size. Parametric tests (One- and Two-Way ANOVA) were used only if data were normally distributed. Otherwise, nonparametric alternatives were chosen (Mann-Whitney and Kruskal-Wallis). Data are presented as mean ± standard error of the mean (SEM). p values ≤ 0.05 were considered significant (*p ≤ 0.05, **p ≤ 0.01, ***p ≤ 0.001, ****p ≤ 0.0001). The specific statistical test used is indicated in each figure legend.

## Acknowledgements

We would like to thank Nathalie Jurisch-Yaksi for helpful and meaningful discussion about our data; Dr. Gislene Pereira for providing LRRC45 antibody and for input on the manuscript; Dr. Mirko Cortese for initial help with EM analysis; Marleen Trapp, Anna-Lena Schramm, Johanna Huber, Alina Rißberger, Aurica L. Ritter and Nico Burmistrak for technical support and producing preliminary data; the Interdisciplinary Neurobehavioral Core (INBC) at Heidelberg University and the German Cancer Research Center (DKFZ) animal facility for animal housing, caring and logistic help; the Genomics and Proteomics Core Facility (DKFZ) for providing the RT-qPCR machine Roche LightCycler480, the Light Microscopy Facility (DKFZ) for providing the cryostats and confocal microscopes. Finally, we thank all Patrizi group members for helpful discussions.

## Authors information

### Authors and Affiliations

**Schaller Research Group, Department of Neuronal Signaling and Morphogenesis, German Cancer Research Center (DKFZ), Heidelberg, Germany**

Kim Hoa Ho, Valentina Scarpetta, Annarita Patrizi

**Faculty of Biosciences, Heidelberg University, Heidelberg, Germany**

Kim Hoa Ho

**Department of Neurosciences Rita Levi Montalcini, University of Turin, Turin, Italy**

Valentina Scarpetta, Marco Sassoè-Pognetto

**Department of Veterinary Sciences, University of Turin, Grugliasco, Italy**

Chiara Salio

**Optical Microscopy Facility, Max-Planck-Institute for Medical Research, Heidelberg, Germany**

Elisa D’Este

**Department of Molecular Neurogenetics, Center for Molecular Neurobiology, ZMNH, University Medical Center Hamburg-Eppendorf, Hamburg, Germany**

Matthias Kneussel

**Abberior Instruments GmbH, Göttingen, Germany**

Martin Meschkat, Christian A. Wurm

**Institut Curie, PSL Research University, Université Paris-Saclay, Orsay, France**

Maria M. Magiera

**Institute of Neuronal Cell Biology, Technical University of Munich, Munich, Germany**

Monika S. Brill

### Authors contribution

K.H.H. and A.P. conceived and designed the study; K.H.H., C.S., M.S.P., M.M.M., M.S.B. and A.P. designed the experiments; K.H.H., V.S., C.S., M.S.B., M.M.M. and A.P. performed experiments; K.H.H. performed snRNA analysis; E.D.E., M.M. and C.A.W. provided STED microscopy data; M.K and J.K. helped with methodology; K.H.H., V.S. and A.P. analyzed data and interpreted the results; K.H.H., M.S.P., M.S.B. and A.P. wrote the manuscript. A.P. supervised the study. All co-authors read, edited, and approved the manuscript.

## Availability of data and materials

All data generated during this study are included in this published article. The developed analysis tools are available from the corresponding author upon request.

