## Supplemental Data for "Intrinsic microtubule destabilization of multiciliated choroid plexus epithelial cells during postnatal lifetime"

### Ho et al., Supplementary Materials

**Supplementary Table 1. Detailed information of postmortem choroid plexus human samples**

| Case n. | Age (years) | Gender | Experimental use | Biobank |
|---|---|---|---|---|
| 1275 | 2 years | F | IF | NIH |
| 4942 | 0 years | F | IF | NIH |
| 5576 | 3 years | M | IF | NIH |
| 5754 | 0 years | M | IF | NIH |
| 5770 | 0 years | F | IF | NIH |
| 5618 | 68 years | F | IF | NIH |
| 5744 | 84 years | F | IF | NIH |
| 5824 | 74 years | M | IF | NIH |

**Supplementary Table 2: qRT-PCR primer sequences**

| Gene | Primer sequence (5' – 3') | Length | Tm | %GC | Product size (bp) | Source |
|---|---|---|---|---|---|---|
| *Ttll1* | GTGTCGCAGTCCACAAAAGAA | 21 | 59.33 | 47.62 | 93 | PrimerBank[1] |
| | ATACAGACGAAGGTCAAACTTCC | 23 | 58.43 | 43.48 | | |
| *Ttll2* | ATGACCAATCTCACCAGGAAGG | 22 | 59.76 | 50.00 | 76 | PrimerBank[1] |
| | AGAGAGACTCACCATAGCGAC | 21 | 58.71 | 52.38 | | |
| *Ttll3* | ATGGGCCGACTCAGAAACG | 19 | 60.08 | 57.89 | 104 | PrimerBank[1] |
| | CGCAAGAGACACCGGATCA | 19 | 59.78 | 57.89 | | |
| *Ttll4* | AGGCAAAAACCATACCAGCAA | 21 | 58.68 | 42.86 | 119 | PrimerBank[1] |
| | GTGGCTTTATTGGCAGTGACA | 21 | 59.11 | 57.62 | | |
| *Ttll6* | AAGCCCTTCATCATCGACGG | 20 | 60.18 | 55.00 | 142 | PrimerBank[1] |
| | TGTCTAGGTTAGGGTGGGAGTAA | 23 | 59.66 | 47.83 | | |
| *Ttll8* | TTCACAACCAAGATTGGACTCTG | 23 | 58.86 | 43.48 | 108 | PrimerBank[1] |
| | GCTCTCAGTGCAAAGGCCATA | 21 | 60.68 | 52.38 | | |
| *Ttll11* | TGTCTCAGGTCCAAACGGTG | 20 | 59.89 | 55.00 | 76 | PrimerBank[1] |
| | CAACCACTATCCGGTTTCACAA | 22 | 59.86 | 45.45 | | |
| *Ttll13* | GTTAGAGAGGAATCTACCACCCC | 23 | 59.36 | 52.17 | 97 | PrimerBank[1] |
| | CTACAACCGATAAGGCAACCC | 21 | 58.71 | 52.38 | | |
| *Hdac6* | GAGGAGCTGATGTTGGTTCAC | 21 | 58.92 | 52.38 | 124 | PrimerBank[1] |
| | AGTTCGGATGCAGATACACTGA | 22 | 59.24 | 45.45 | | |
| *Sirt2* | CAAGGAAAAGACAGGCCAGA | 20 | 57.72 | 50.00 | 84 | Valle et al., 2014[2] |
| | GCCTTCTTGGAGTCAAAATCC | 21 | 57.14 | 47.62 | | |
| *Aurka* | GAATTGTAACGGCTGAGCTACC | 22 | 59.39 | 50.00 | 73 | Designed by the author |
| | TCCATGTCACAGGCAGGAAC | 20 | 59.96 | 55.00 | | |
| *Agbl5* | CTGCTCATTCTCGTCTTCAGG | 21 | 58.46 | 52.38 | 101 | PrimerBank[1] |
| | ATCGAGTCCTAATGCAAGGGA | 21 | 58.6 | 47.62 | | |
| *Agtpbp1* | ATGAGCAAGCTAAAAGTGGTGG | 22 | 59.18 | 45.45 | 107 | PrimerBank[1] |
| | TCTGACTCTGTGGAATCCGTATT | 23 | 58.98 | 43.38 | | |
| *Agbl2* | ATGAATGTCCTGCTTGAGATGG | 22 | 58.45 | 45.45 | 146 | PrimerBank[1] |
| | CAAACGCGCTGATGAGTGC | 19 | 60.51 | 57.89 | | |
| *Agbl3* | TGACTTGGATGAGGATTCCTTCA | 23 | 59.15 | 43.48 | 96 | PrimerBank[1] |

| | | | | | | |
|---|---|---|---|---|---|---|
| | GGGAAGAATGGGTCACCAATAG | 22 | 58.44 | 50.00 | | |
| *Agbl4* | AGGCAGGCAATGATACAGGAA | 21 | 59.43 | 47.62 | 135 | PrimerBank[1] |
| | GGTTACCACTTTCAAAGCAAGCA | 23 | 60.18 | 43.48 | | |
| *Atat1* | AGCAACCGGCACGTTATTTAC | 21 | 59.53 | 47.62 | 147 | PrimerBank[1] |
| | GCAAAGGGGTTCTACCTCATTGT | 23 | 60.81 | 47.83 | | |
| *Gas8* | GACGAGATGAACGAGATGCAG | 21 | 58.88 | 52.38 | 120 | PrimerBank[1] |
| | TTCCCATTTCAGGTCCTTTAGC | 22 | 58.3 | 45.45 | | |
| *Rsph9* | CACTGCTCACGTCCCTTATGC | 21 | 61.34 | 57.14 | 100 | PrimerBank[1] |
| | GCGATGTAGTAATCCGCCACA | 21 | 60.54 | 52.38 | | |
| *Cfap53* | TCTAGTTTCCTGTCGCCACC | 20 | 59.39 | 55.00 | 134 | Designed by the author |
| | CTCTGAGCTTTGGGAGGCTT | 20 | 59.67 | 55.00 | | |
| *Ccdc40* | GGAAGGCCCTCCGAGGAATA | 20 | 60.76 | 52.38 | 86 | PrimerBank[1] |
| | TTGGGGTCTGACCTTTCTGTG | 21 | 59.86 | 52.38 | | |
| *Dnah9* | CACAGTGAGCCAGTCTCAGAT | 21 | 59.45 | 52.38 | 144 | Designed by the author |
| | GGGGGTGGGATTTTGTCCTT | 20 | 59.88 | 55.00 | | |
| *Hydin* | CTTGCCCCTCCGAATCAGAG | 20 | 60.18 | 60.00 | 114 | PrimerBank[1] |
| | AGGATTGCCTCGTAACAGTGA | 21 | 59.1 | 47.62 | | |
| *Kif3a* | ATGCCGATCAATAAGTCGGAGA | 22 | 59.37 | 45.45 | 134 | Designed by the author |
| | GTTCCCCTCATTTCATCCACG | 21 | 58.98 | 52.38 | | |
| *Spast* | GCCTCGATCAAAAACTGTCCT | 21 | 58.57 | 47.62 | 106 | PrimerBank[1] |
| | TCCCGGTCTTGCTCCAGAAA | 20 | 61.13 | 55.00 | | |
| *Hprt* | CAAACTTTGCTTTCCCTGGT | 20 | 56.72 | 45.00 | 101 | Ho and Patrizi, 2021[3] |
| | TCTGGCCTGTATCCAACACTTC | 22 | 60.03 | 50.00 | | |
| *Rpl13a* | AGCCTACCAGAAAGTTTGCTTAC | 23 | 58.93 | 43.48 | 129 | Ho and Patrizi, 2021[3] |
| | GCTTCTTCTTCCGATAGTGCATC | 23 | 59.51 | 47.83 | | |
| *Rpl27* | AAGCCGTCATCGTGAAGAACA | 21 | 60.27 | 47.62 | 143 | Ho and Patrizi, 2021[3] |
| | CTTGATCTTGGATCGCTTGGC | 21 | 59.67 | 52.28 | | |

**Extended Figure 1. Choroid plexus anlage only contains undifferentiated and unciliated epithelial cells. a**, Immunofluorescence images of telencephalic (t) choroid plexus (ChP) at E12.5. Sagittal brain sections were stained with antibodies against E-cadherin (yellow) and Ki67 (white). Inset boxes feature ChP proximate (1) or distant (2) from the ventricular wall, respectively. **b**, Immunofluorescence images of hindbrain (h) and tChP at E11.5 and E12.5 respectively. Sagittal brain sections were stained with antibodies against $\gamma$-Tubulin ($\gamma$-Tub, green) and acetylated tubulin (AcTub, magenta). Inset boxes feature ChP adjacent (1) or distant (2) from the ventricular wall, respectively. Scale bars: overview: 100 $\mu$m, inset: 10 $\mu$m.

**Extended Figure 2. The proliferation of the hindbrain and telencephalic choroid plexus epithelial cells follows similar trajectory**. **a**, E12.5 hindbrain (h) choroid plexus (ChP) sections were stained with antibodies against EdU (magenta) and BrdU (green). Insets feature ChP close (1) or distant (2) from the ventricular wall, respectively. Scale bar: overview: 100 $\mu$m, inset: 10 $\mu$m. **b,** Quantification of the percentage of BrdU-positive (+) choroid plexus epithelial cells at four embryonic time points in both h- and telencephalic (t) ChP. *n* = 3 mice. Bars represent Mean ± SEM.

**Extended Figure 3. Embryonic choroid plexus contains differentiated monociliated epithelial cells. a,** Immunofluorescence images of telencephalic (t) choroid plexus (ChP) at E16.5 of Otx2-GFP mice. **b,** Immunofluorescence images of E16.5 choroid plexus epithelial cells close to the ventricular wall (Root). ChP sections were stained with antibodies against Ki67 (white) and glutamylated tubulin (GluTub, magenta). Asterisks (*) display Otx2-GFP cells negative for Ki67 and positive for GluTub. **c,** Transmission electron microscopy (TEM) serial reconstruction showing monociliated cells at E16.5 ChP. Arrows indicate outer dynein arms. **d,** Quantification of different type of ciliated stages at E16.5 and P0 ChP. *n* = 3 mice per age. Two-Way ANOVA, $^{**}p \leq 0.01$, $^{****}p \leq 0.0001$. Bars represent mean ± SEM. Scale bars: 100 $\mu$m (**a**); 5 $\mu$m (**b**); overview 1 $\mu$m, inset 200 nm (**c**).

**Extended Figure 4. Ciliogenesis progress for each ventricle. a,** Immunofluorescence images of E16.5 (left) and P0 (right) choroid plexus (ChP). ChP sections were stained with antibodies against $\gamma$-Tubulin ($\gamma$-Tub, green) and acetylated tubulin (AcTub, magenta). Arrows indicate cilia orientation towards cerebrospinal fluid (CSF). Scale bar: 5 $\mu$m. **b,** Schematic representation of orientation score. **c,** Quantification of ciliated cells with a selective orientation. *n* = 3 mice.

Contingency Chi-squared test. **d-f,** Quantification of the percentages of cells positive for multiple basal bodies (**d**), percentages of total ciliated cells (**e**) and cilia length (**f**) at E14.5, E16.5 and E18.5 in telencephalic (t) and hindbrain (h)ChP. Bars represent mean ± SEM, *n* = 3 mice per age (**d, e**); Median ± 95% CI, *n* = 130-200 cilia (from 3 mice) per ventricle per age (**f**). Two-Way ANOVA, *p≤0.05, **p≤0.01, ***p≤0.001, ****p≤0.0001. ns: not significant.

**Extended Figure 5. Acetylation, glutamylation and glycylation cover the whole length of ChP cilia's axonemes.** Immunofluorescence images of choroid plexus (ChP). **a-c,** ChP sections were stained with antibodies against Tubulin alpha 4a (Tub4a, green) and acetylated tubulin (AcTub, magenta) (**a**); glutamylated tubulin (GluTub, green) and AcTub (magenta) (**b**); glycylated tubulin (GlyTub, white) and GluTub (green). Dotted line indicates region of interest showing GluTub-positive/GlyTub-negative cilia at E16.5. Scale bar: 3 μm (**a, c**).

**Extended Figure 6. Sustained basal body localization of CFAP53 at postnatal ages. a,** Absolute RT-qPCR expression level of a subset of motile cilia genes in E16.5 choroid plexus (ChP) homogenates. *n* = 3 embryos. **b,** Diagram of an axonemal cross section of canonical motile cilia configuration and identified selected genes. **c,** Immunofluorescence images of ChP. P35 ChP sections stained with antibodies against GluTub (green) and CFAP53 (magenta). Scale bars: 5 μm.

**Extended Figure 7. Disappearance of choroid plexus cilia occurs in both ventricles with a similar dynamic. a-c,** Quantification of the percentage of cells positive for multiple basal bodies (**a**), percentages of total ciliated cells (**b**) and cilia length (**c**) at multiple postnatal ages in both hindbrain (h) and telencephalic (t) ChP. Two-Way ANOVA, *p≤0.05, **p≤0.01, ***p≤0.001, ****p≤0.0001. Bars represent Mean ± SEM, *n* = 3 mice per age (**a, b**); Median ± 95% CI, *n* = 80-200 cilia (from 3 mice) per ventricle per age (**c**). **d,** Heatmap of snRNA-seq data of differentially expressed cilia organization genes in all three ventricles over 3 developmental ages. dChP: diencephalic choroid plexus.

**Extended Figure 8. Uncharacteristic short length of choroid plexus axonemes. a, b,** Transmission electron microscopy (TEM) images showing sagittal cross sections of multiple type of axonemes. **b,** TEM images showing multiple cilia retreated deep into cytoplasm and surrounded by intracellular vacuoles. Scale bars: 1 μm (**a**); 200 nm (**b**). **c,** Quantification of axoneme length distribution in adult choroid plexus epithelial cells. Bars represent Median ± 95% *n* = 205 axonemes, 3 mice.

**Extended Figure 9. Expression profiles of posttranslational modification catalyzing enzymes and the tubulin-severing enzyme Spastin across ages. a, b,** Pseudo-heatmap of RT-qPCR expression profiles of acetylases, glutamylases, glycylases (**a**) and deacetylases and deglutamylases (**b**) across ages (from E16.5 to P720). **c,** Relative mRNA levels of Spastin from E16.5 to P720. *n* = 3 mice. Bars represent Mean ± SEM, One-way ANOVA test. ns: not significant.

**Extended Figure 10. Characterization of human choroid plexus epithelial cells. a,** Immunofluorescence images of human postmortem choroid plexus (ChP). ChP sections were stained with antibodies against Transthyretin (TTR, red), OTX2 (yellow) and E-cadherin (E-cad, green). Scale bars: 20 μm. **b,** Quantification of the percentage of cells positive for multiple basal bodies in young patients (0-3 years-old) and aged patients (60–90 years-old). Bars represent Mean ± SEM, *n* = 5 young specimens, 3 aged specimens. Mann-Whitney test, ns, not significant.

Extended Figure 1

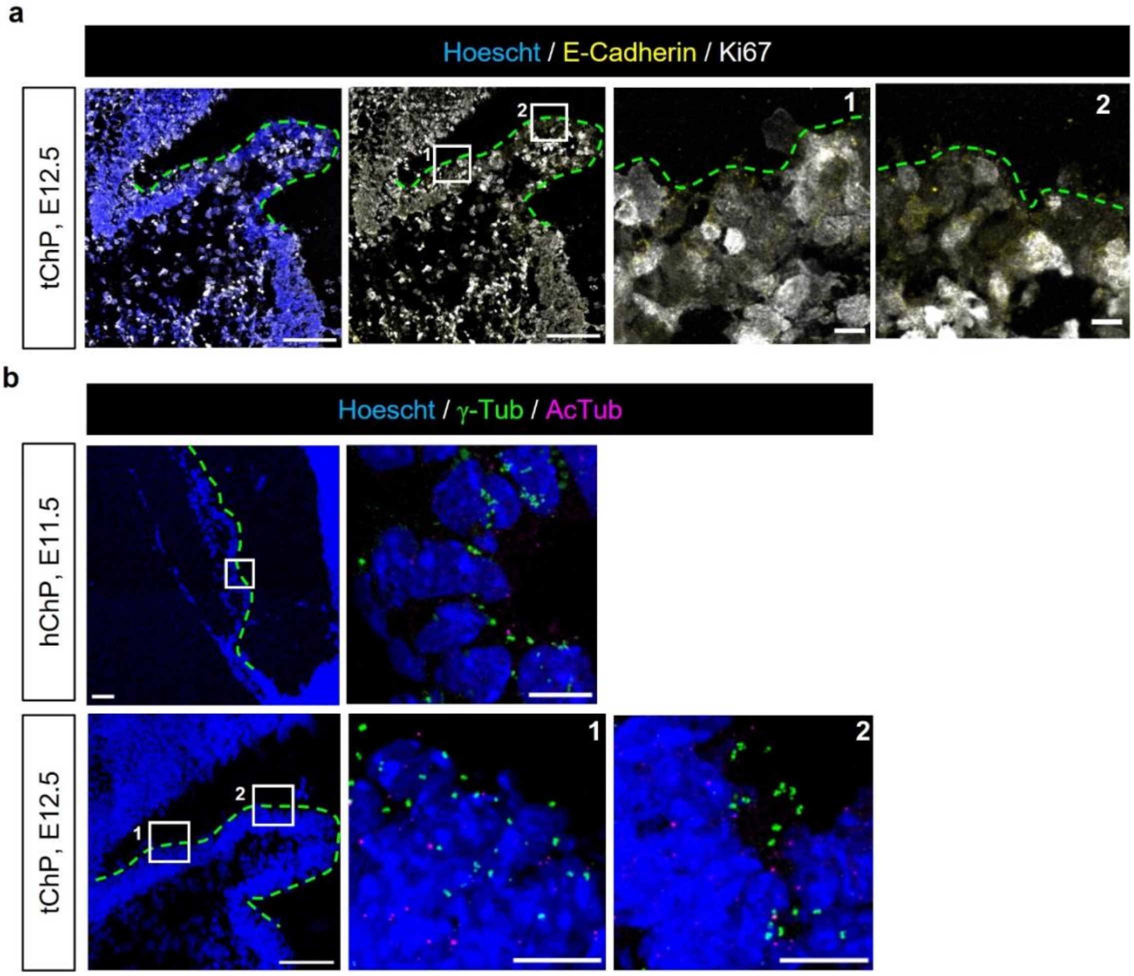

Extended Figure 2

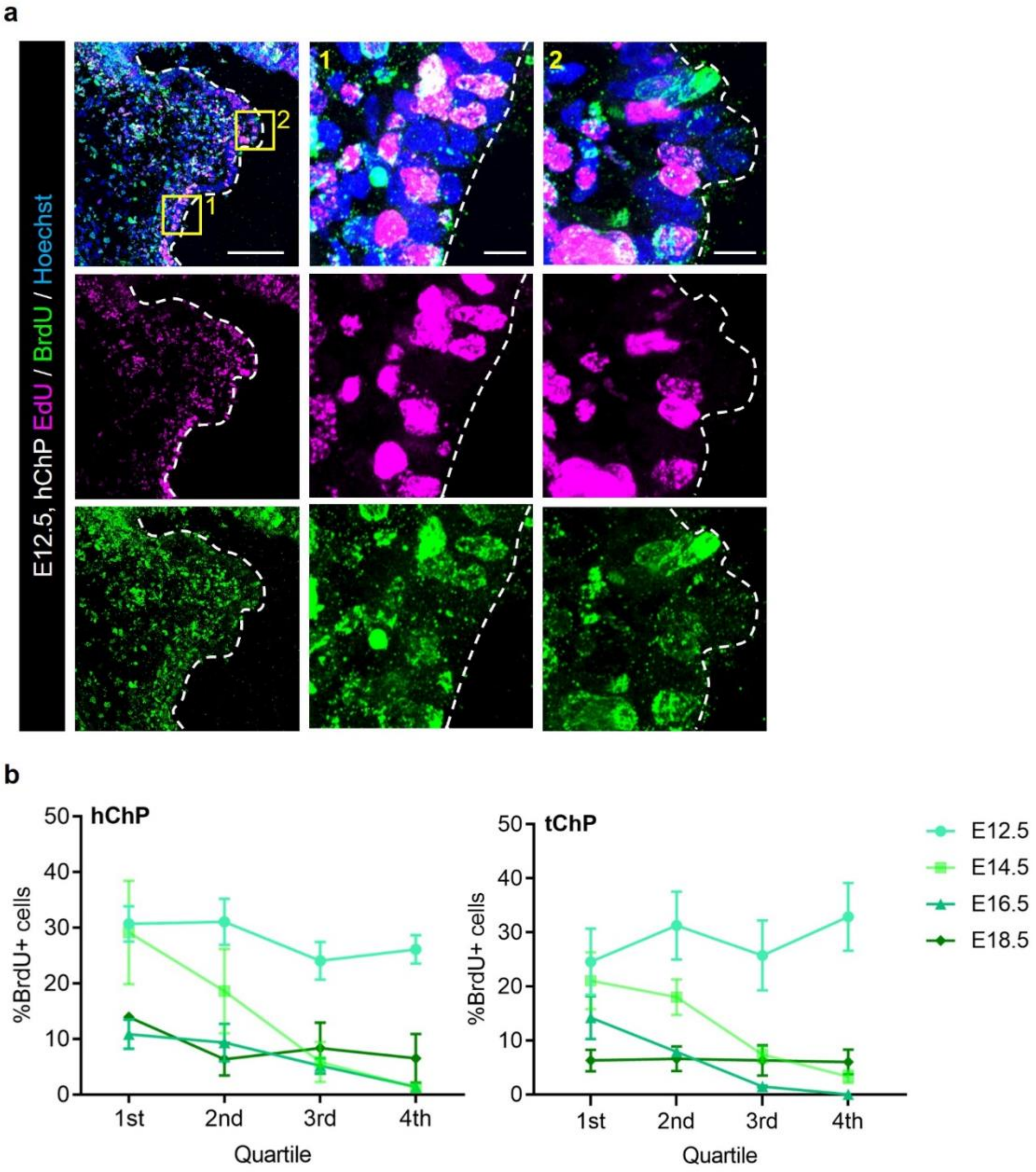

Extended Figure 3

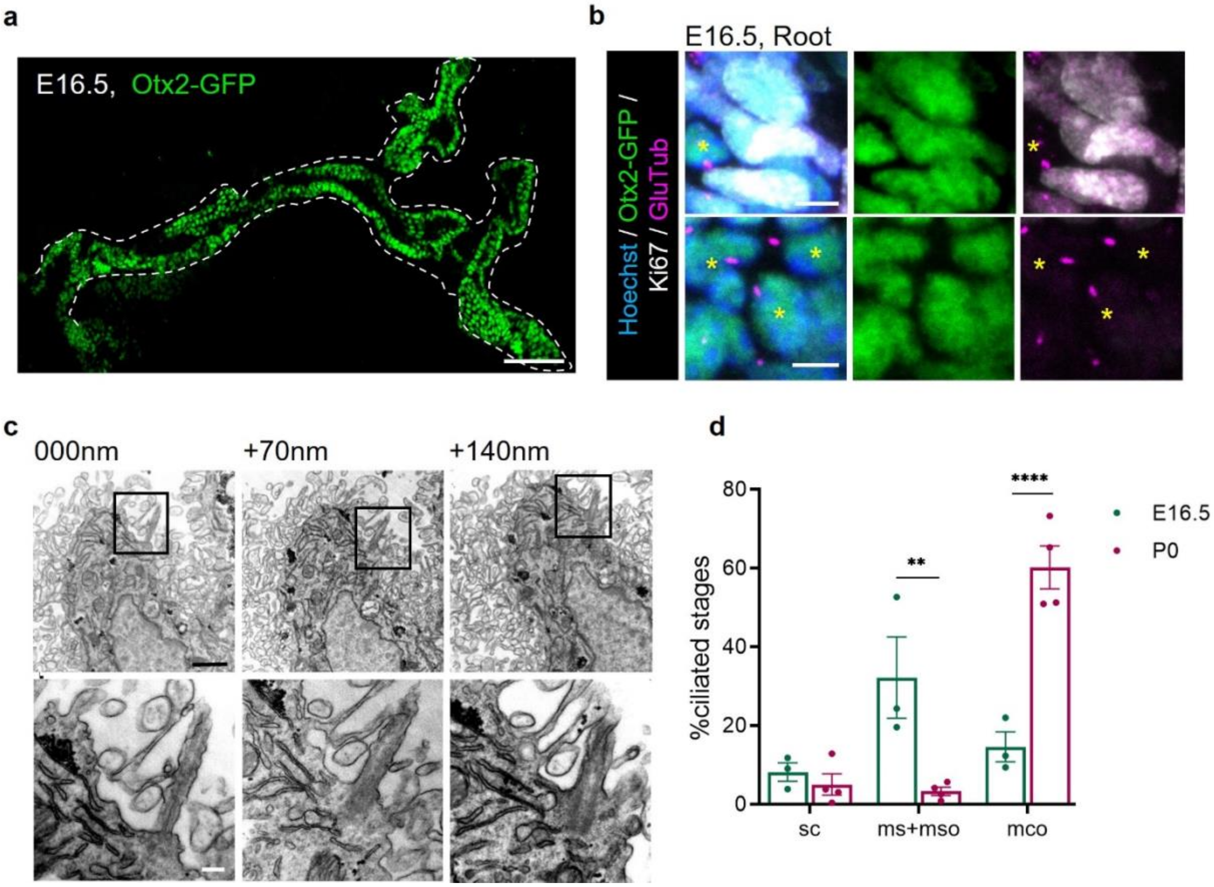

Extended Figure 4

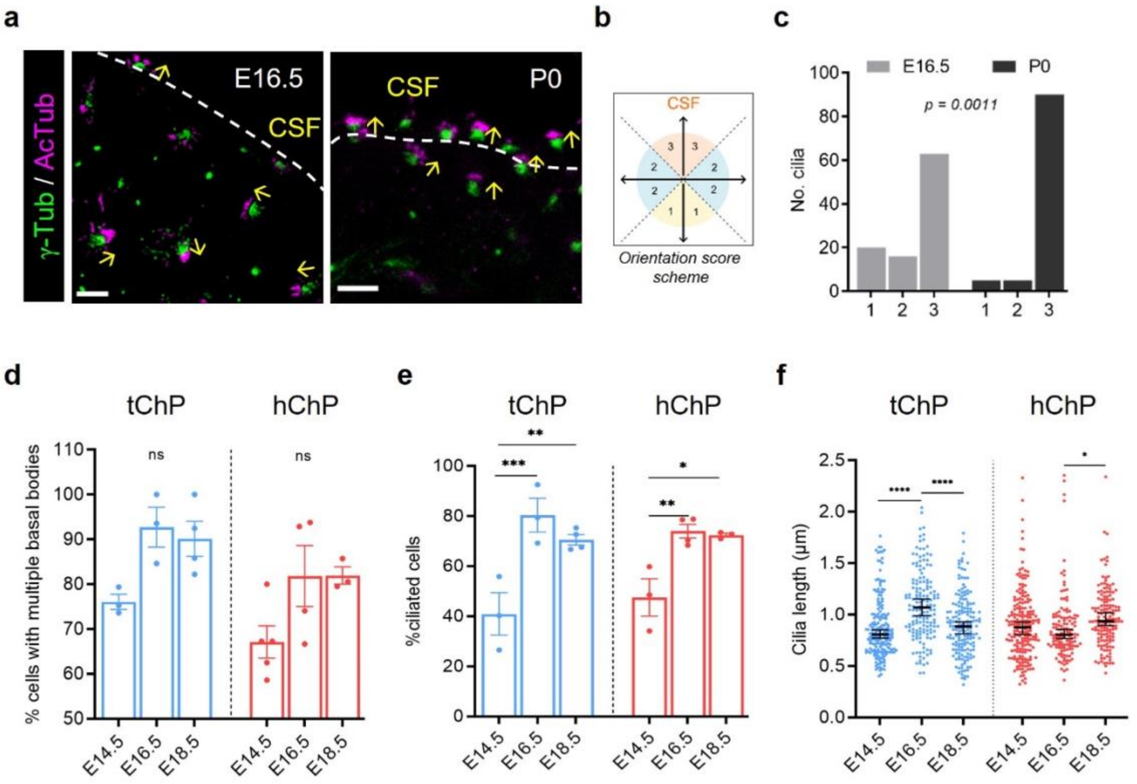

Extended Figure 5

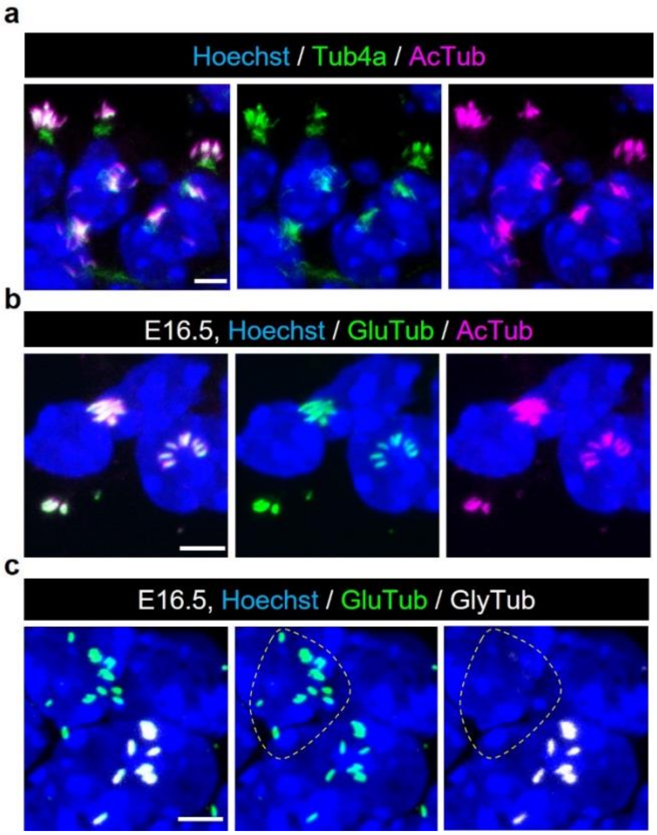

Extended Figure 6

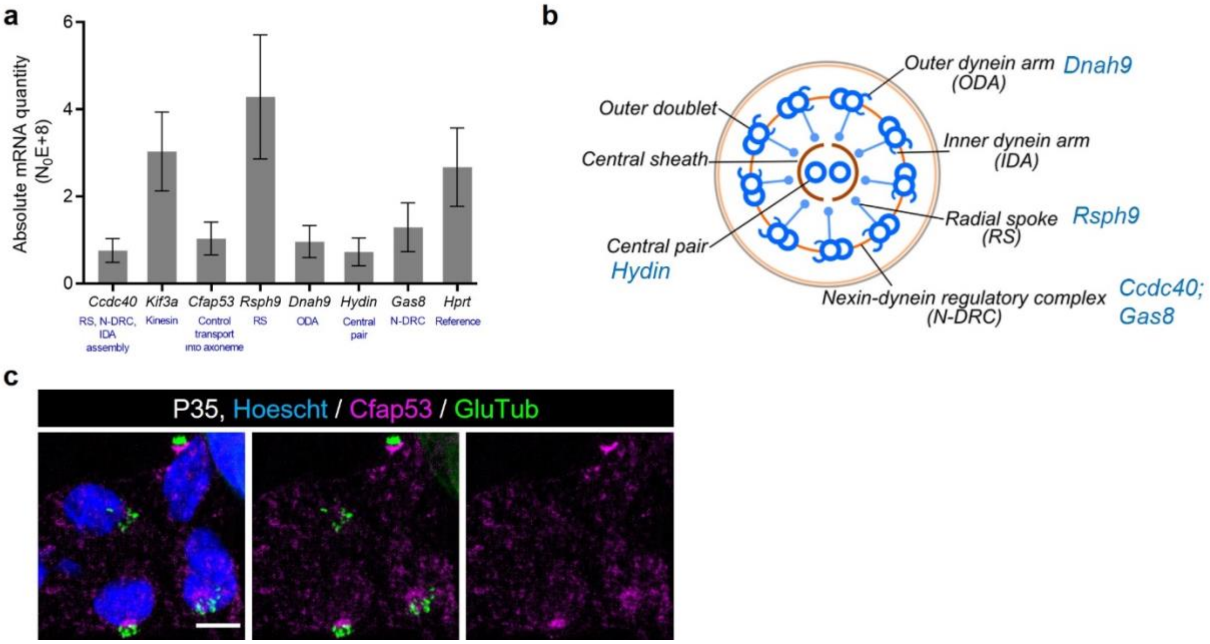

Extended Figure 7

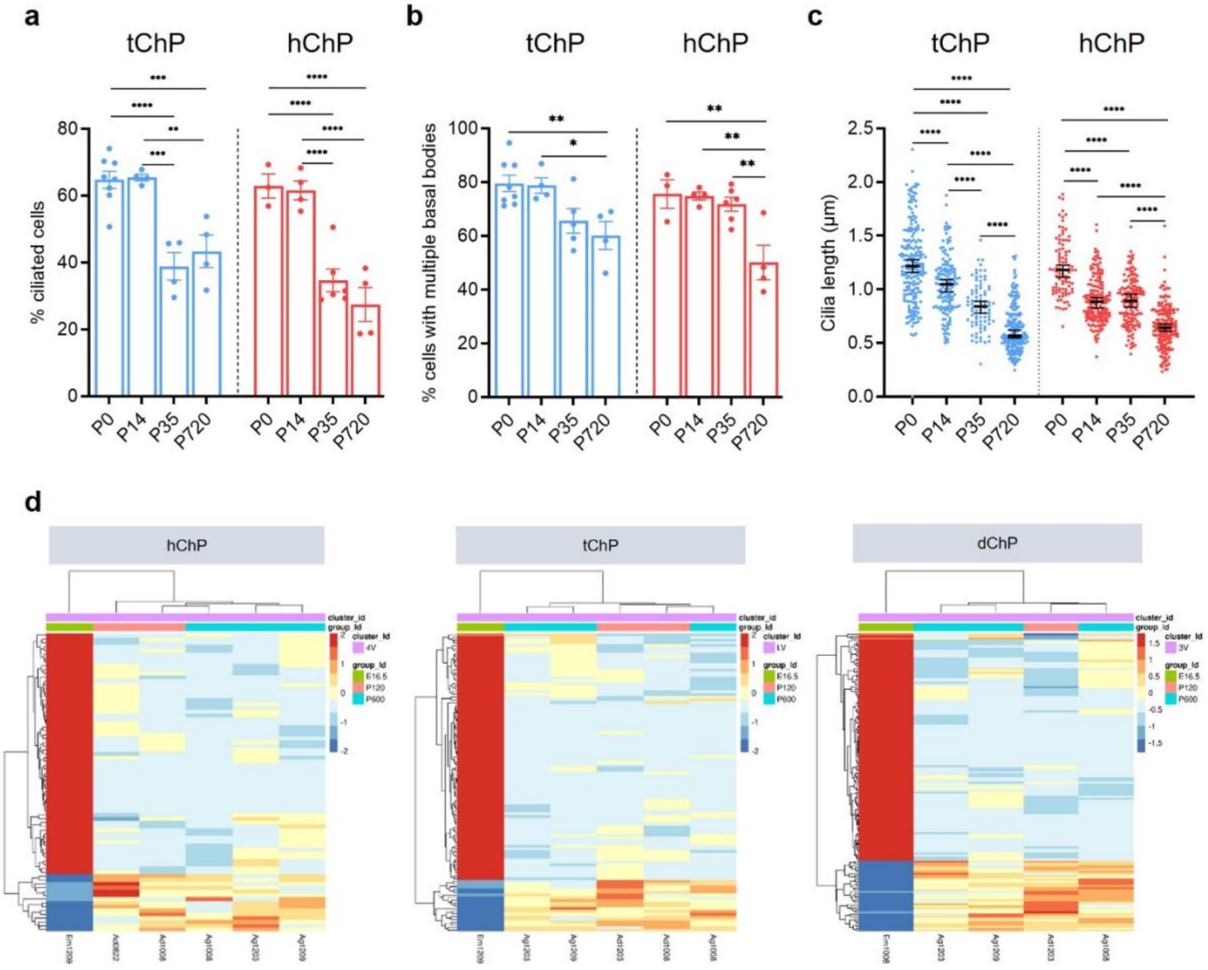

Extended Figure 8

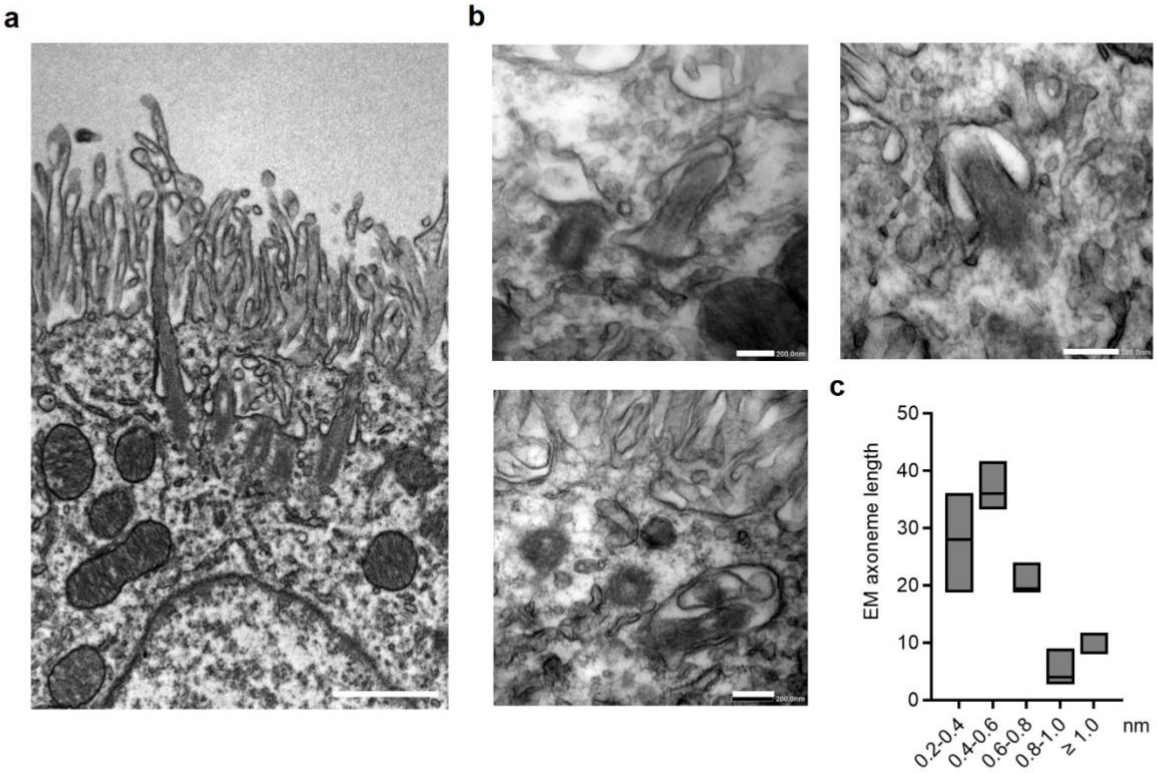

Extended Figure 9

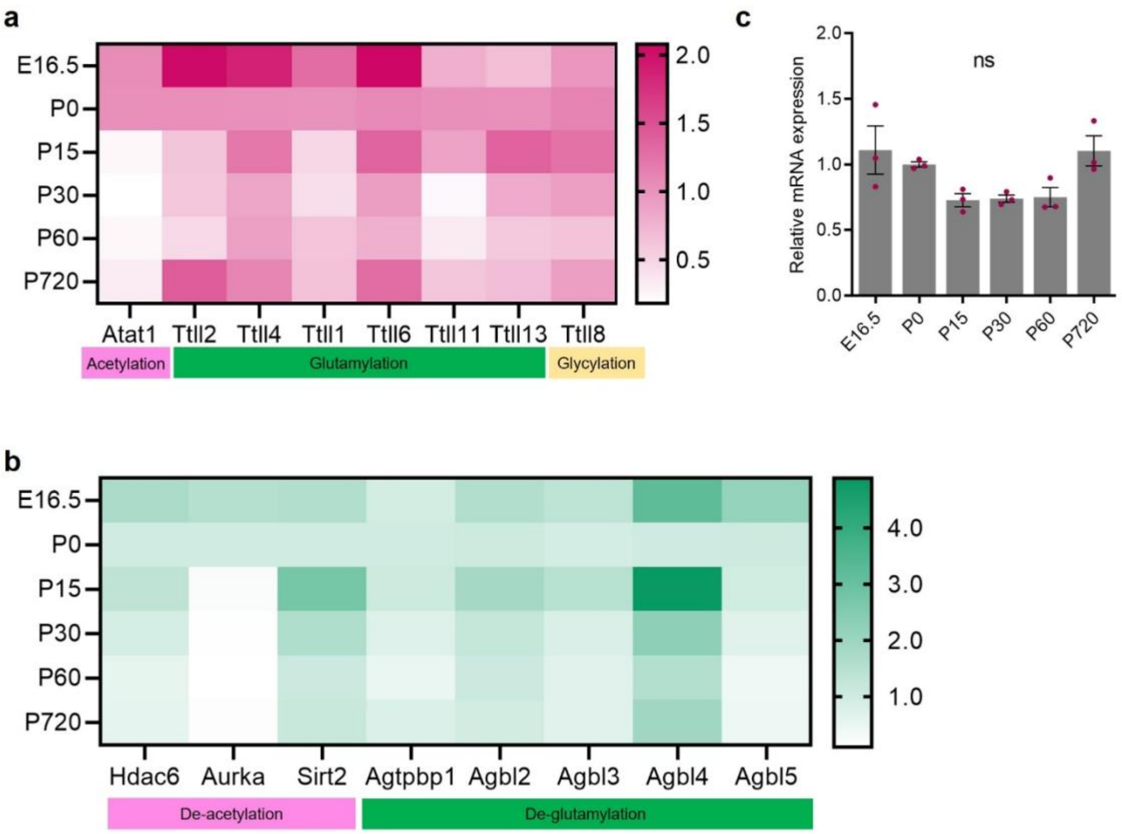

Extended Figure 10

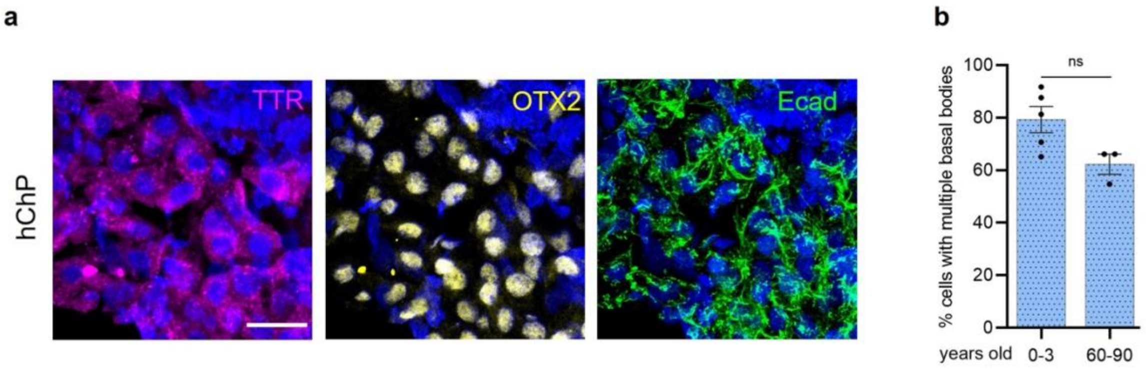
